# Optimization and scaling of patient-derived brain organoids uncovers deep phenotypes of disease

**DOI:** 10.1101/2020.08.26.251611

**Authors:** Kevan Shah, Rishi Bedi, Alex Rogozhnikov, Pavan Ramkumar, Zhixiang Tong, Brian Rash, Morgan Stanton, Jordan Sorokin, Cagsar Apaydin, Anthony Batarse, Julia Bergamaschi, Robert Blattner, Spencer Brown, Anthony Bosshardt, Carlos Castrillo, Brenda Dang, Shiron Drusinsky, Luigi Enriquez, David Grayson, Juliana Hilliard, Pei-Ken Hsu, Chili Johnson, Ryan Jones, Andy Lash, Chia-Yao Lee, Kelly Li, Austin McKay, Elliot Mount, Justin Nicola, Ismael Oumzil, Justin Paek, Deborah Pascoe, Arden Piepho, Sean Poust, Daphne Quang, Matthew Schultz, Jessica Sims, Patrick Taylor, Geffen Treiman, Oliver Wueseke, Noah Young, Alex Pollen, Doug Flanzer, Daniel Chao, Gaia Skibinski, Saul Kato, G. Sean Escola

**Affiliations:** System1 Biosciences, Inc; Zuckerman Institute, Department of Psychiatry, Columbia University; Weill Institute for Neurosciences, University of California, San Francisco

## Abstract

Cerebral organoids provide unparalleled access to human brain development in vitro. However, variability induced by current culture methodologies precludes using organoids as robust disease models. To address this, we developed an automated Organoid Culture and Assay (ORCA) system to support longitudinal unbiased phenotyping of organoids at scale across multiple patient lines. We then characterized organoid variability using novel machine learning methods and found that the contribution of donor, clone, and batch is significant and remarkably consistent over gene expression, morphology, and cell-type composition. Next, we performed multi-factorial protocol optimization, producing a directed forebrain protocol compatible with 96-well culture that exhibits low variability while preserving tissue complexity. Finally, we used ORCA to study tuberous sclerosis, a disease with known genetics but poorly representative animal models. For the first time, we report highly reproducible early morphological and molecular signatures of disease in heterozygous TSC+/− forebrain organoids, demonstrating the benefit of a scaled organoid system for phenotype discovery in human disease models.

---

Recapitulation of brain development with three-dimensional (3D) culture of induced pluripotent stem cells (iPSCs) has ushered in a new era of human systems neurobiology. Patient-derived organoids (PDOs) exhibit the cellular diversity and early organization of specific human brain areas, functional neural network activity^1–3^, and the genetics of the affected individual^4^. Despite these promising advances, the variability of organoid culture remains a major challenge for reliable use in disease modeling, especially when simultaneously studying multiple donors^5^.

Most published organoid protocols, which require pooled culture in bioreactors, spinner flasks, or larger cell culture plates^1,2,6,7^, limit understanding of PDO variability for several reasons. First, these culture formats are incompatible with automation equipment like liquid handlers, necessitating the use of manual processes. Manual culture is extremely time consuming, hindering replication over factors that influence phenotypic results such as the donor from which an iPSC line was derived (“donor”), the specific iPSC clone from which an organoid was seeded (“clone”), and the group of organoids seeded at the same time (“batch”) **(Fig. 1a)**. Without characterizing donor, clone, and batch effects, the risk of reporting artifactual phenotypes increases significantly. Second, manual culture exacerbates variations due to human-introduced inconsistent liquid transfers and organoid handling. Third, cross-laboratory reproducibility of results is a major concern in 3D culture, as small experimental differences lead to significantly different outcomes^8–10^. Fourth, pooled culture exacerbates batch effects^11^, possibly due to inter-organoid signaling. Finally, pooled culture prevents assaying of individual organoids at multiple time points, presenting inherent limitations on multimodal/multitemporal disease phenotyping.

**Fig. 1.**
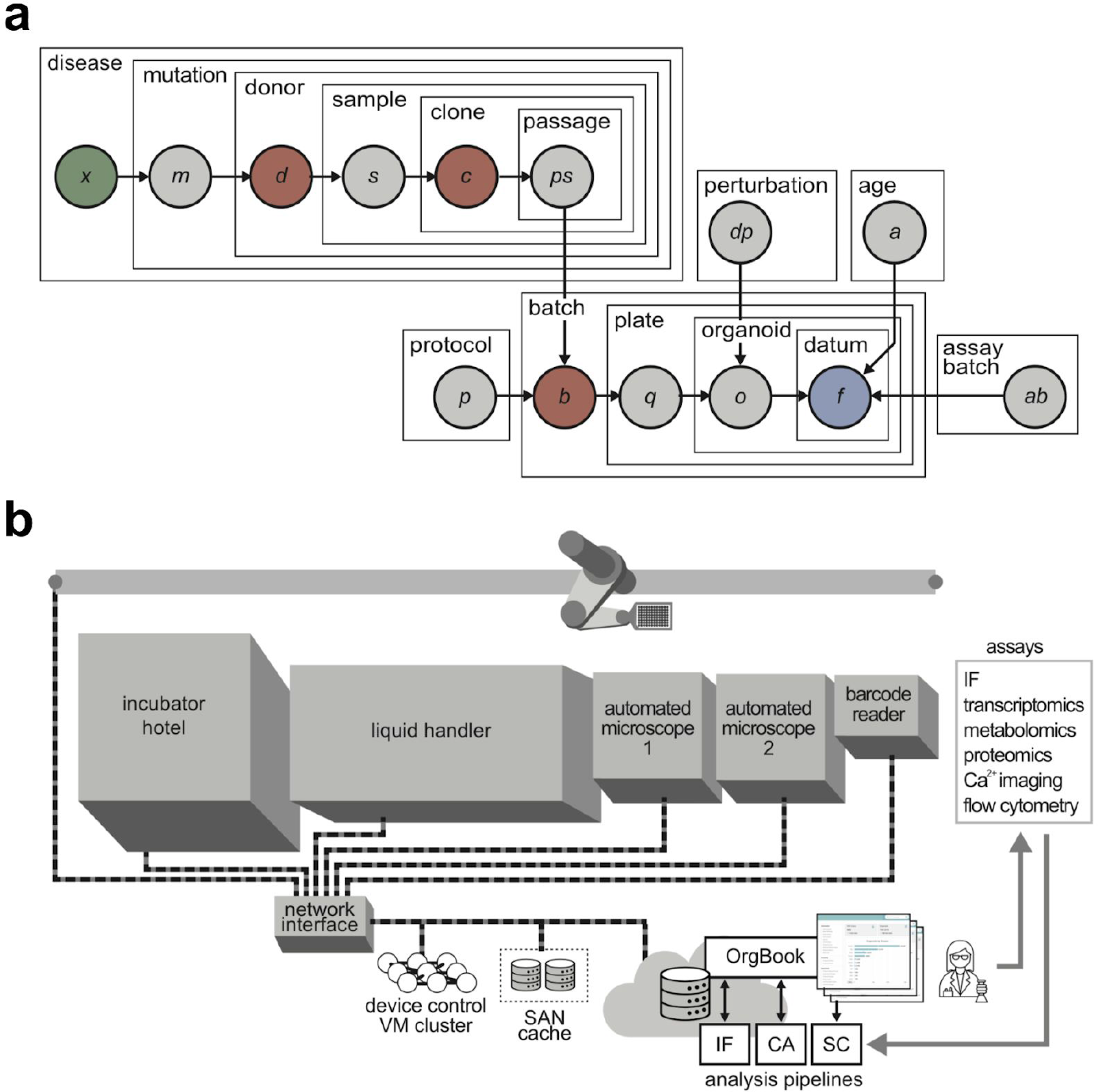
Sources of variability and ORCA system overview. (**a**) The dependence hierarchy of identified potential sources of variability in data collected from a particular organoid. The objective of deep phenotyping is to correctly attribute signal in observed data (blue) to disease state (green), in spite of confounding variability introduced by biological and technical sources. In this work we focus on the influence of three major confounds: donor, clone, and batch (red). (**b**) Schematic overview of the Organoid Culture and Assay (ORCA) platform. Experiments designed with cloud-based custom software (Orgbook) are scheduled and executed on a local virtual machine (VM) cluster that controls the robotic systems to perform various activities such as feeding, imaging and assaying. Data generated from ORCA and metadata from each event (e.g., metadata about a feeding event, or the outputs of an assay) are cached on a local storage area network (SAN), then uploaded to a cloud database and staged for automated analysis and data visualization pipelines (e.g., immunofluorescence (IF), calcium imaging (CA), and single-cell RNA-seq (SC)).

Addressing the challenge of organoid variability, we present: (1) an integrated Organoid Culture and Assay platform (ORCA) that enables standardized and automated culture conditions, scalable individual-well organoid culture **(Fig. 1b; Supplementary Fig. 1),** (2) a novel machine-learning enabled method for characterizing organoid variability: the “rank-to-group score”, and (3) a directed forebrain differentiation protocol (DFP) compatible with automated longitudinal 96-well plate organoid culture.

Using the DFP and ORCA, we investigated tuberous sclerosis (TSC), a genetic disorder that causes benign tumors in the skin, kidneys, lung, and brain^12^. 90% of patients have debilitating neurological symptoms including epilepsy, autism spectrum disorder, and cognitive impairment^13,14^. Mutations in the TSC1 or TSC2 genes account for 85% of TSC patients^15^. Loss of TSC1 or TSC2 in rodents leads to increased cell size, altered cell morphology, and subependymal hamartomas^16,17^, but fails to induce cortical tubers, defined by macroscopic regions of disorganization and dysfunction^18^. Thus, better, ideally human, models are needed for investigating TSC and potential therapeutics^16,17^.

Previous human 3D TSC models required long-term culture of organoids past day 100^19^, or biallelic inactivation (i.e., TSC2^−/−^) to reveal disease phenotypes^20^. However, only one-third of patient samples exhibit biallelic inactivation^21^. Here, using deep phenotyping – an unbiased, multimodal phenotype discovery approach – we report highly reproducible phenotypes in heterozygous TSC^+/−^ organoids at culture day 35, including machine learning-derived morphological signatures via immunofluorescence and cell-type specific differential gene expression via single-cell RNA-sequencing. These results point the way towards enhanced disease biology characterization and therapeutics for TSC.

## RESULTS

### ORCA enables longitudinal tracking of individual complex forebrain organoids

ORCA cultures organoids in individual wells (24- or 96-well formats) using automated liquid handlers for aspirating, feeding, media sampling, and harvesting (**Fig. 1b**), eliminating variability introduced by human organoid handling. ORCA integrates automation hardware control and Orgbook, our software tracking, experiment design, and automated analysis platform (**Supplementary Fig. 1**), reducing bookkeeping error and human labor.

ORCA-cultured organoids maintain rich biological characteristics: self-organization into radially-structured neuroepithelial buds^22^ (Day 30; **Fig. 2a**) with diverse cellular composition^3,23^ (Day 90; **Fig. 2b**) and endogenous neural network activity^6,11^ (Day 90; **Fig. 2f**). Well-formed, Sox2+ neuroepithelial buds show robust Emx1 expression (Day 30; **Fig. 2a, top**) and Tbr2+ subventricular zone (SVZ) development with cortical Emx1+ and Ctip2+ neurons accumulating at the surface of the organoid (**Supplementary Fig. 2**). Organoids show cortical plate (CP) formation: Satb2+ layer 4 and Tbr1+ layer 6/subplate (Day 80; **Fig. 2a, bottom**). Organoids exhibit cellular compositions of excitatory neurons, inhibitory neurons, astrocytes, and radial glia (Day 90; **Fig. 2b**).

**Fig. 2.**
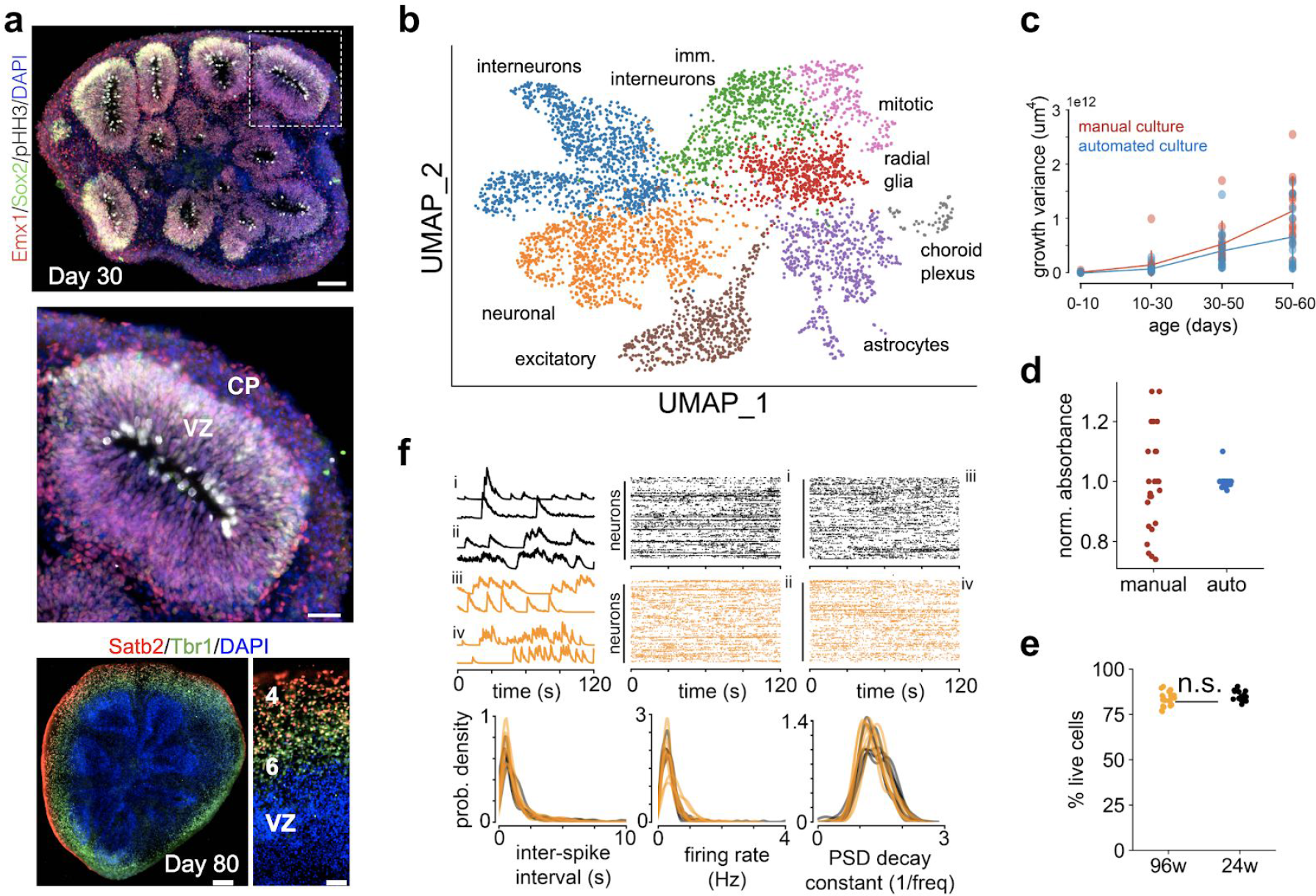
Organoids grown in automated culture display consistent hallmarks of brain development. **(a)** Top: Immunofluorescence image of day 30 DFP organoid; scale bar is 100 μm. Middle: Zoom-in of a bud with clear emergence of cortical plate (CP) and ventricular zone (VZ); scale bar is 30 μm. Bottom: Immunofluorescence image of day 80 DFP organoid; scale bars are 200 μm (bottom left) and 50 μm (bottom right). CP: cortical plate; VZ: ventricular zone; 4: layer 4; 6: layer 6. **(b)** scRNA-seq derived gene expression patterns for cells from day 90 DFP organoids shown in the first two UMAP dimensions and colored by cell type (6 donors; 1 cell pool per donor; 3 organoids per cell pool). **(c)** Variance in growth curves during four age ranges (estimated via spline-regression, see methods). Automated culture reduces variability compared to manual culture for later growth (d50-d60) (2-way ANOVA; p<0.001; model DF = 7) **(d)** The volume of media in each well after manual feeding (left: CV: 17% over 24 wells) is more variable as measured by plate reader absorbance (phenol red at 560 nm) compared with automated feeding (right: CV: 2% over 24 wells). **(e)** Percent of live cells at day 91 measured by flow cytometry over DFP organoids grown from 3 iPSC clones (each data point is an organoid). The 96w group (orange) was kept in 96w plates throughout its entire lifetime. The 24w group (black) were grown in 96w plates until day 20 and then transferred to 24w plates for the rest of their lifetime. **(f)** Top left: calcium time series of two representative neurons from four different DFP organoids (i-iv). Black traces are organoids grown in 24w conditions while orange traces are those grown in 96w conditions (as in **e**). Top right: rasters of inferred spikes from the same four videos (i-iv). Bottom: distributions of derived statistics from the inferred neural spikes demonstrate similar neural dynamics between 24w and 96w conditions.

Compared with manual culture, organoids grown using ORCA exhibit more stereotyped growth dynamics, as measured by reduced size variance (**Fig. 2c**; *p* < 0.001; 2-way ANOVA), likely resulting from increased reliability in hitherto manual processes. For example, ORCA reduced the variation in the final volume of media in our culture wells post-feeding (coefficient of variation 2% with automated aspiration, vs. 17% manual) (**Fig. 2d**).

ORCA facilitated scale-up of individual well culture to increase replication. Compared with manual methods, media exchange increased from 20 to 72 24-well plates per person hour (with 4 automated feeders running in parallel), enabling the simultaneous culture of over 40,000 individually tracked organoids. ORCA also enabled scale up of assay throughput. Quantitative PCR by manual processing (involving sample collection, RNA extraction, quantification, cDNA synthesis and qPCR) took 33 hours per 100 samples; after automation, 100 samples took 5 hours, and resulted in a reduction in variance across replicates (97.2% of technical replicates showed less than one fold change in cycle threshold or *C_t_* after automation vs. 86.3% before) (**Supplementary Fig. 3**).

We used ORCA to test indefinite culture in 96 well plates versus transfer to 24 well plates at day 20. After compensating for reduced media volume in 96 well plates with more frequent feeding, we found no effect on organoid health as measured by the percentage of live cells following dissociation (**Fig. 2e**). Moreover, we found no differences in the neural dynamics of the maturing organoids between the 24-well and 96-well protocols (**Fig. 2f**). Thus, we conclude that 96-well culture does not compromise the viability or developmental trajectory of cortical organoids, while allowing four times as many organoids in the same spatial footprint.

As discussed next, longitudinal tracking of single organoids enables (1) measurement and characterization of confounding sources of variability across multiple assay modalities simultaneously, and (2) rapid and systematic exploration of organoid protocols in the quest for reduced variance across donors, clones, and batches. Together, these open the door to disease phenotyping with patient-derived organoids.

### Variability is correlated across phenotypic modalities and attributable to known experimental variables

The goal of phenotyping is to identify signatures across multiple modalities that map specifically to disease and not to other confounds that may influence organoid development. We decomposed the variability in our unpatterned protocol (UP) organoids with respect to a set of known biological and technical confounds including: the *donor* from which an iPSC cell line was derived, the specific iPSC *clone* from which an organoid was seeded (multiple iPSC clones may be derived from a single donor), and the *batch* of organoids seeded at the same time. (Each batch comprised organoids cultured in individual wells from multiple clones using the same reagents; **Supplementary Fig. 4**.)

Low-magnification brightfield imaging is both non-destructive and easily automatable, making it valuable for quality control and phenotyping. However, it is not obvious *a priori* that gross organoid morphology would correspond to meaningful differentiation features. We found that the presence of small outgrowths on the perimeter of organoids corresponds to high WNT1 expression – a caudal marker of neural tube and a caudalizing morphogen^24,25^ – versus a smoother exterior on the perimeter of the organoids with low WNT1 (**Fig. 3a**). To systematically interrogate the relationship between brightfield morphology and differentiation outcome, we developed a machine learning method capable of learning low-dimensional representations, or vector embeddings, of organoid images (**Fig. 3b**). Specifically, we trained a convolutional neural network to discriminate between self – images taken of the same organoid, including at different ages and orientations – and other – images of different organoids. We then extracted the vector embeddings learned to solve this discrimination task and found that they correlated remarkably well with gene expression of multiple developmental markers measured at day 14 (**Fig. 3c**). Therefore, brightfield imaging provides biologically meaningful information, establishing it as a scalable, non-destructive assay for organoid protocol development and quality control.

**Fig. 3.**
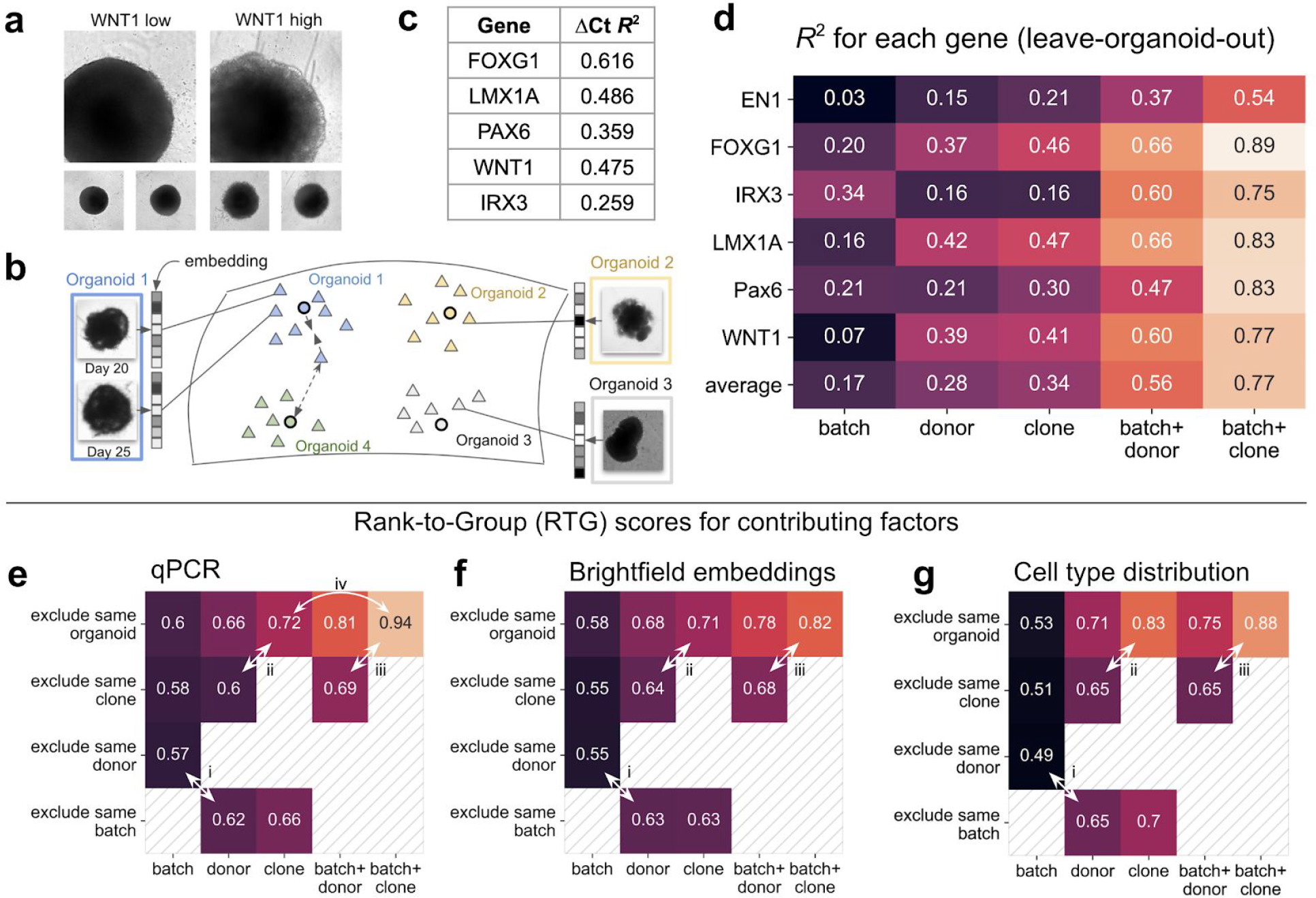
Donor, clone, and batch contributions to variability correlate across phenotypic modalities. **(a)** Organoid brightfield images at day 14 with maximal and minimal WNT1 expressions as measured by qPCR relative to endogenous control B2M. **(b)** Schematic representation of the machine learning model trained to extract useful vector embeddings from organoid brightfield images. The model is trained to cluster vector embeddings of images of the same organoid more closely to their own center than to cluster centers of any other organoids (as shown by dashed arrows).These learned vector embeddings can be used for various tasks, such as correlating to gene expression (as in C) or estimating the contribution of different sources of variability to brightfield morphology (as in F). **(c)** Correlation of learned brightfield embeddings to gene expression of key developmental marker genes (both brightfield and qPCR at day 14), estimated by predicting gene expression values using brightfield embeddings with regularized regression, in a leave-one-organoid-out cross-validation scheme. **(d)** Effect of each property (or set of properties) in each column on the expression of the gene in each row. Each value shown is R^2^ computed using only that property/set of properties. Value of 1 indicates that the given property/set of properties perfectly predicts the expression of the given gene. **(e, f, g)** Estimates of how much different sources of variability contribute to observed phenotypic features by rank-to-group (RTG) score. A value of 1 (maximum possible) indicates that all organoids most similar to a query organoid share the same property/set of properties as the query organoid; while a value of 0.5 (minimum possible) indicates that organoids sharing a property/set of properties are no more similar to the query organoid than would be expected by chance. Arrows highlight specific comparisons – i: batch vs. donor; ii: donor vs. clone; iii: batch+donor vs batch+clone; iv: batch+clone vs clone. RTG scores determined for **(e)** gene expression using qPCR, **(f)** brightfield morphology vector embeddings, and **(g)** cell type distributions identified with scRNA-seq.

Next, we partitioned variability into its hierarchical sources, both within and across assays. Specifically, we modeled qPCR gene expression as a function of donor, clone, batch, and their interactions, and observed for several genes that donor effects dominate batch effects (average *R*^2^ of 0.28 vs. 0.17, respectively). Thus, batch variability is sufficiently low to detect the differences between donors (**Fig. 3d**).

To assess donor, clone, and batch effects across assays, we developed a novel statistical method, the rank-to-group (RTG) score, which evaluates contribution of confounding group (e.g., donor, clone, or batch) membership to any assay-derived features (e.g., gene expression values, brightfield vector embeddings, or cell type proportions). The RTG score quantifies the extent to which similar organoids according to a particular assay share the same confounding group (**Supplementary Fig. 5**).

For gene expression, the importance of donor and batch evaluated using the RTG score agrees with the results in Fig. 3d (**Fig. 3e-i,** RTG score of 0.62 vs. 0.57, respectively). Additionally, the overall pattern of contribution of each confounding variable is mirrored between the regression results and the RTG scores (each row in **Fig. 3d** vs top row in **Fig. 3e)**.

Clone contributes additional information beyond donor (**Fig. 3e-ii**, RTG score 0.72 vs. 0.6, respectively), suggesting that variability from iPSC reprogramming and/or culture substantially affects differentiation outcomes even for clones sharing a common genetic background, in agreement with a recent study^26^. For this result, the contribution of clone is determined by comparing each organoid to all other organoids, while the contribution of donor is estimated by comparing only to organoids from different clones. This “exclude-same-clone” evaluation strategy is important because it ensures that the RTG score does not attribute variability to donor that is more appropriately explained by clone.

We also interrogated the effect of donor and clone within a given batch. The interaction between batch and clone explains more variability than the interaction between batch and donor (**Fig. 3e-iii**, RTG score 0.94 vs. 0.69, respectively), further confirming that the particular clone contributes to variability beyond the donor source. Furthermore, it clarifies the role of batch in organoid variability: while batch alone is a weak predictor of gene expression, individual clones are significantly more consistent within a batch than across batches (**Fig. 3e-iv**, RTG score 0.94 vs. 0.72, respectively).

We repeated our RTG score analysis for brightfield vector embeddings (**Fig. 3f**) and cell type distributions derived from scRNA-seq (**Fig. 3g**), and found that the relative importance of donor, clone, batch, and their interactions mirrors gene expression. In all cases, donor explains more variability than batch (**Figs. 3e, f, g-i**, RTG score of 0.62 vs. 0.57, 0.63 vs. 0.55, and 0.65 vs. 0.49, respectively), while clone explains additional variability beyond donor (**Figs. 3e, f, g-ii**, RTG score of 0.72 vs. 0.6, 0.71 vs. 0.64, 0.83 vs. 0.65 respectively). Finally, for all 3 assays, organoids that share batch and clone are highly stereotyped, more so than organoids that only share batch and donor (**Figs. 3e, f, g-iii**, RTG score of 0.94 vs. 0.69, 0.82 vs. 0.68, and 0.88 vs. 0.65, respectively).

That variability assessments from three distinct assays (qPCR at day 14, brightfield imaging at day 14, and scRNA-seq cell types at day 35) are in strong agreement (average pairwise Spearman’s rank correlation of 0.924) supports using cheap, non-destructive modalities like brightfield imaging to understand the contributions of confounds like donor, clone, and batch across assays.

In summary: first, donor and clone have greater influence than batch; second, organoids from the same clone are more similar than organoids that merely share the same donor; third, while batch identity is only a weak predictor of organoid features, organoids sharing the same batch and donor are much more similar than organoids sharing just the same donor; and finally, organoids sharing the same batch and clone are much more similar than organoids sharing just the same clone. These results have implications for well-powered experimental design and statistical interpretation of results: generalizability across datasets comprising several donors, clones and batches is a must for robust disease phenotyping.

### ORCA identified a directed forebrain-specific protocol with reduced inter-organoid variability

Disease phenotyping requires reproducible organoid culture from a variety of iPSC cell lines with varying genetic backgrounds. Directed protocols mitigate donor-to-donor variability by preserving cellular and structural complexity while minimizing undesirable differentiation trajectories^3,27–29^. Although prior efforts have produced directed protocols that improve organoid consistency, generalization of the protocols to support many donor lines requires systematic multi-parameter optimization. This requires a scaled system and detailed analysis of variability for rapid iteration. We applied the ORCA platform as a protocol development engine for simultaneous optimization over multiple parameters (**Supplementary Table 1**), starting from the unpatterned protocol (UP) analyzed in Fig. 3. This led to a directed forebrain protocol (DFP) compatible with indefinite single-well culture that robustly produces forebrain tissue over a large panel of donors (**Supplementary Table 1**).

We grew organoids from 58 iPSC clones derived from 21 donors in individual wells and assayed their variability. Using qPCR at day 14, we found that FOXG1 expression – a key marker of forebrain induction – varied over five orders of magnitude in the UP protocol (**Fig. 4a**; colorbar). Furthermore, these organoids exhibited high variability across the developmental genetic program, as visualized by principal components analysis (PCA) for four other key developmental genes (Wnt1, LMX1A, Pax6, and IRX3) (**Fig. 4a**). Increased FOXG1 expression is correlated with decreased variability among these genes as evidenced by a tight cluster for FOXG1-high organoids (brighter color), indicating that FOXG1 expression is a useful surrogate for more consistently patterned organoids across genes. Compared with UP organoids, FOXG1 expression measured at day 14 in DFP organoids has a much higher mean and a much lower variance (ΔC_t_ relative to housekeeping gene GAPDH: mean ± stdev; UP: −9.00 ± 4.97; DFP: −1.69 ± 1.46; *p* < 1e–50, two-sided *t*-test; **Fig. 4b**).

**Fig. 4.**
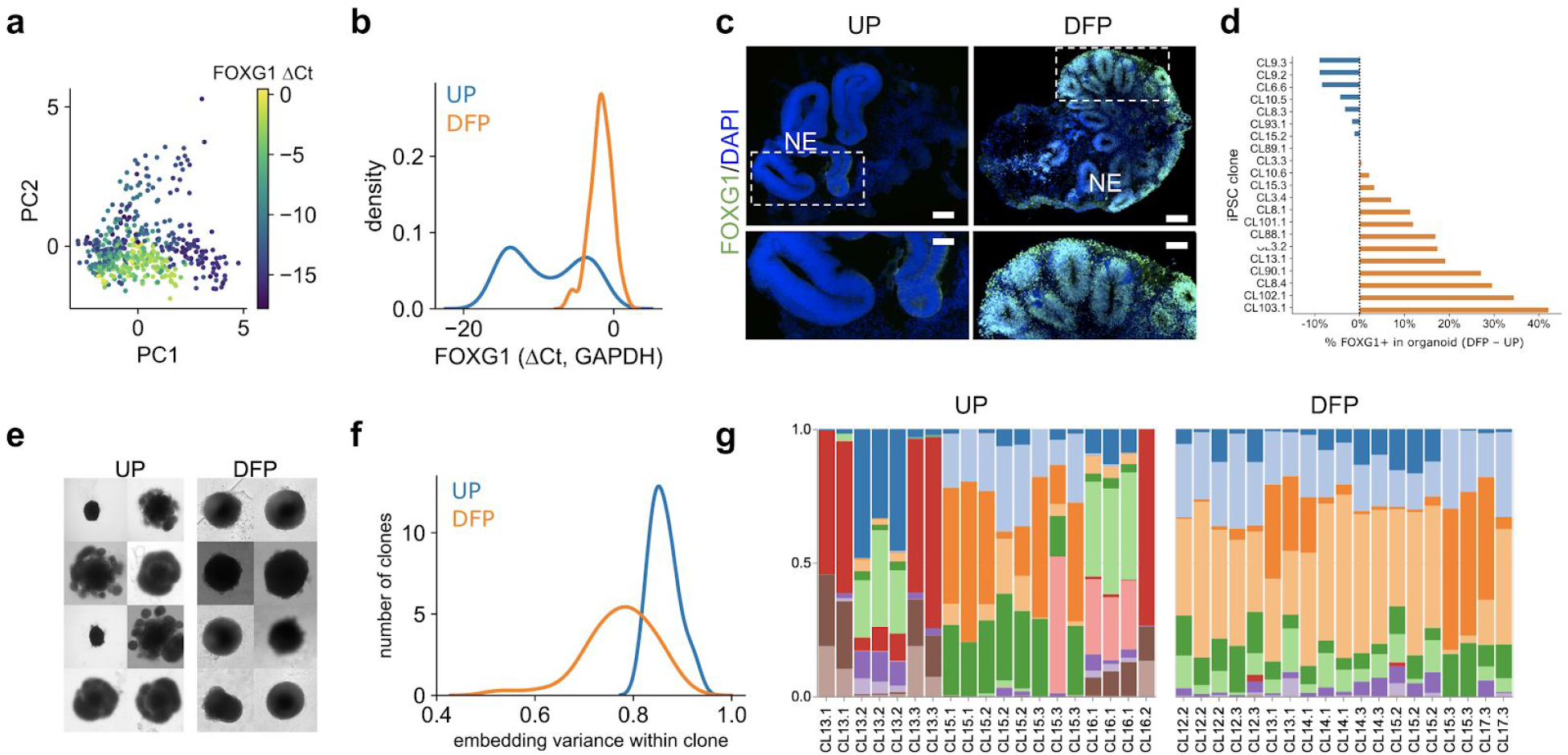
DFP organoids show reduced variability in morphology, gene and protein expression, and cellular composition. **(a)** Principal component analysis computed on Wnt1, LMX1A, Pax6, IRX3 gene expression in UP organoids, as measured by qPCR and colored by FOXG1 expression (ΔCt w.r.t. GAPDH). Each point is an organoid. **(b)** Distribution of log-fold change (ΔCt w.r.t. GAPDH) values of FOXG1 over 486 UP organoids (representing 58 clones derived from 21 distinct donors, grown over 17 batches (average of 12 clones represented per batch)); and 149 DFP organoids (representing 54 clones derived from 20 distinct donors, grown over 6 batches (average of 11 clones represented per batch)). **(c)** Representative immunofluorescence images stained for FOXG1 at day 35 of the same clone cultured using UP (left) or DFP (right). Insets are high magnification images of boxed regions. Scale bar: 120 μm; 60 μm in insets. NE: neuroepithelium. **(d)** Comparison of total FOXG1 immunofluorescence at day 35 over all clones grown using both protocols. **(e)** Grid of selected organoids demonstrating brightfield UP and DFP organoid morphologies imaged at day 21, from the same clone. **(f)** Histogram of intra-clone variance of brightfield vector embeddings for organoids grown using UP and DFP. **(g)** Relative distribution of cell types measured by scRNA-seq for day 35 UP organoids (3 unique donors, 8 unique clones) compared with DFP organoids (6 unique donors, 8 unique clones). Each column is a unique organoid and each color represents a computationally identified cell type cluster. Colors are matched across UP and DFP subplots.

Additionally, using immunofluorescence imaging at day 35, UP organoids frequently exhibited bud-like structures that were FOXG1-negative (**Fig. 4c**). In contrast, FOXG1 positivity increased by 48% in DFP organoids. FOXG1^+^ buds indicate that forebrain-specific cells are self-organizing into neuroepithelial structures^22,30^. DFP organoids demonstrated significantly greater FOXG1 expression than UP organoids across clones (*p* < 2e-22, ANOVA) (**Fig. 4d**). The decreased variance and increased mean of FOXG1 expression across both mRNA and protein assays demonstrate the success of ORCA-enabled search for a robust forebrain protocol.

Brightfield imaging at day 21 for UP organoids showed gross structural heterogeneity (**Fig. 4e**). The same clones grown using the DFP, however, showed reduced intra-clone variability as measured by the variance of vector embeddings derived with the same machine learning method described in **Fig. 3b** (*p* < 2e-8, Kolmogorov–Smirnov test, **Fig. 4f**). Images of more clones grown using the DFP exhibit similar homogeneity **(Supplementary Fig. 6)**.

The variability in developmental fate rescued by the DFP extends to the cellular composition of the organoid. We performed scRNA-seq on 19 UP and 20 DFP day 35 organoids from a subset of the same clones and clustered the cells’ gene expression patterns, allowing us to identify putative cell types. We observed significant cell type heterogeneity across organoids and cell lines in UP organoids (**Fig. 4g**, left), corroborating previously reported results^3^. Notably, entire cell types present in some organoids are absent in others. In contrast, DFP organoids exhibited a marked reduction in variability (**Fig. 4g**, right), also replicating previous reports about reduced cell type variability in directed protocols^3^.

In summary, we observe reductions in variance and improvements in desired forebrain specification over four different assay modalities: brightfield imaging, qPCR, scRNA-seq, and immunofluorescence imaging. These multimodal results increase confidence in our directed differentiation protocol and highlight the strength of a scaled, individualized culture approach.

### Growing organoids from many donors reveals reproducible phenotypes of tuberous sclerosis

With the increased sensitivity afforded by low-variability DFP organoids, we applied ORCA to characterize the monogenic neurodevelopmental disease tuberous sclerosis (TSC) known for its hallmark morphological signatures in humans. Since only one-third of clinical cortical tuber samples exhibit biallelic inactivation^21^, we reasoned that monoallelic inactivation should induce aspects of tuberous sclerosis pathology, despite prior reports in organoids finding no pathology. To investigate this, we grew DFP organoids from five TSC^+/−^ heterozygous donors (one TSC1^+/−^and four TSC2^+/−^) (**Supplementary Table 2**), eight unrelated healthy donors, and five donors with a neurodevelopmental disorder unrelated to TSC. We then cryo-sectioned and DAPI-stained day 35 organoids grown from each of these donors to look for morphological signatures of disease.

We developed a machine learning model to classify small (32 μm x 32 μm) patches of DAPI-stained cryo-sectioned organoid images as TSC^+/−^ versus control. The model comprises two stages: an unsupervised feature extractor followed by a supervised classifier (**Fig. 5a**). Like the brightfield model described in Fig. 3, the feature extractor produces a vector embedding for each patch by solving a self-versus-other discrimination task (**Supplementary Fig. 7**). These embeddings are then fed into a classifier which learns to predict whether a patch was from a TSC or control organoid. This two-stage approach has two major advantages. First, the unsupervised nature of the feature extractor enables use of a large training set consisting of immunofluorescence data (674 distinct z-plane regions of tissue sections from 87 organoid sections) not explicitly collected for this TSC experiment. Second, the feature extractor can be explicitly trained to be robust to variations in staining or imaging parameters like focus plane and time between staining and imaging.

**Fig. 5.**
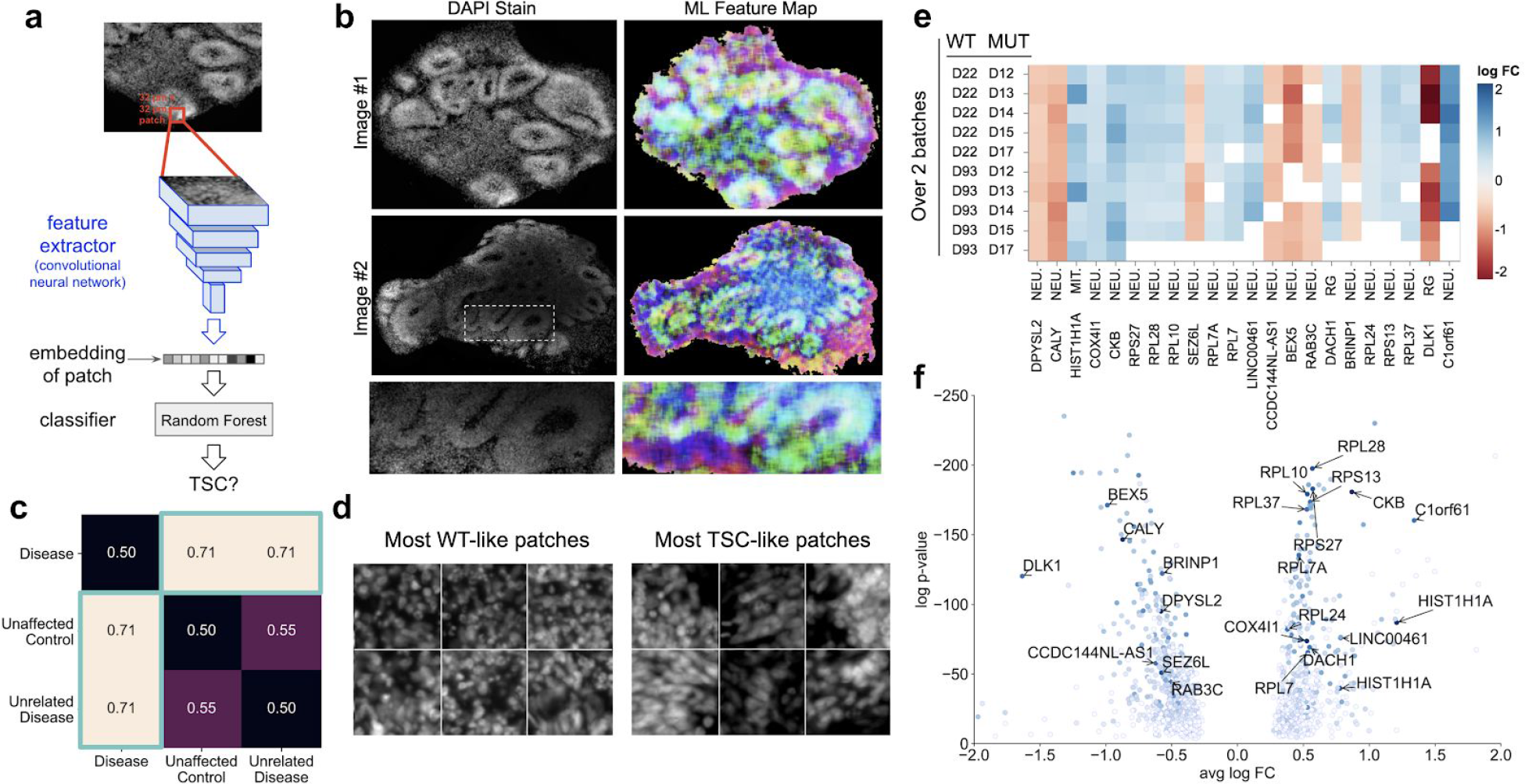
Heterozygous tuberous sclerosis organoids display morphological and gene expression phenotypes. (**a**) Schematic of machine learning system used to classify small patches (red) (32μm x 32μm) as selected from either a TSC sample or a WT sample. (**b**) Left: raw DAPI immunofluorescence images; right: each pixel colored based on the vector embedding of a patch centered on said pixel. More similar colors correspond to nearby vector embeddings, interpretable as similar tissue type. (**c**) Performance of classifier trained on learned vector embeddings to predict TSC disease state. 0.5 indicates no better than random chance. Evaluation performed using leave-one-donor-out cross-validation (AUROC). (**d**) Exemplar patches that scored most highly as WT (top) and those that scored most highly as TSC (bottom). (**e**) DEGs in clusters that are most reproducible over all the 10 possible donor comparisons ((between 5 TSC+/− donors and 2 unrelated healthy donors) separately over two batches of organoids), ordered from left to right by the fraction out of 10 in which the DEG was found in the same cluster. All DEGs shown here were consistent over both batches. A white square indicates the DEG in that cluster was significant in either 0 or 1 batch. Log FC > 0 indicates the gene was upregulated in the TSC organoids. Clusters are abbreviated as follows – NEU: excitatory neurons; MIT: mitotic cells; RG: radial glia. (**f**) Average log2 fold change (x axis) vs. Bonferroni-adjusted p-value (y axis) for all DEGs reproducible in at least 2 out of 10 donor pairs in both batches. DEGs consistent in at least 8/10 donor pairs (the same set as in (e)) are highlighted. The color of each point indicates the fraction out of 10 in which it is significant (darkest blue: 10/10).

We found that learned vector embeddings are interpretable representations of tissue patches. By mapping the principal components of patch embeddings to color, we found that patches from macroscopic features like buds share similar colors across different parts of the same image, and across different images, despite the spatial resolution of a patch being significantly smaller than the size of a macroscopic feature like a bud (**Fig. 5b)**. We were able to robustly classify patches derived from TSC organoid slices on unseen donors versus patches derived from wildtype (WT) organoid slices (leave-donor-out AUROC 0.71; compared to random chance AUROC of 0.5) (**Fig. 5c**). This performance on classifying individual patches translates to perfect performance in predicting whether organoids from a given donor are from a TSC donor or a WT donor (i.e., AUROC 1). Crucially, classifier performance is evaluated using a leave-one-donor-out cross-validation scheme, ensuring that classification performance is not a consequence of irrelevant donor-to-donor morphological differences, but instead of features that are common to TSC organoids of differing genetic backgrounds. Furthermore, the performance of the classifier is as strong when TSC organoids are compared with organoids from MECP2-deficient donors, an unrelated neurodevelopmental disorder (AUROC 0.71 over patches; and perfect performance at the clone-level). This comparison provides evidence that the model is not identifying spurious differences between donors or features of a generic “disease state”, but instead is leveraging features that are specific to TSC. Interestingly, patches that the model scores as most likely to be from a TSC organoid exhibit altered nuclear morphology and organization compared to the most likely WT patches (**Fig. 5d**).

We then performed single-cell RNA sequencing on organoids grown from iPSCs derived from the same five tuberous sclerosis donors and from two unrelated healthy control lines. Each iPSC line was seeded in two separate batches to mitigate the effects of batch variability (**Fig. 5e**). This experimental design enabled the use of stringent cross-batch and cross-donor reproducibility criteria to identify disease-relevant differentially expressed genes (DEGs). We assessed statistically significant differential expression (Bonferroni adjusted p-value < 0.05) above an effect size threshold (|average log_2_ fold change| > 0.25), over *all* 20 possible mutant vs. wild-type comparisons (5 mutant donors x 2 wild-type donors x 2 batches). With these stringent criteria, we identified a set of five robust DEGs, primarily in a cluster of excitatory neurons (enriched for NEUROD6, GPM6A, EMX2, TBR1; **Supplementary Table 3**; **Fig. 5e**).

Two of these five DEGs are downregulated: DPYSL2 (coding for dihydropyrimidinase-related protein 2) and CALY (coding for neuron-specific vesicular protein calcyon). DPYSL2 has been identified to be regulated by mTOR, which is in turn regulated by TSC1/2^31,32^. Functional variants of DPYSL2 are also associated with other neurological disorders - schizophrenia^32^ and de novo autism spectrum disorder^33^ – suggesting a locus of shared pathophysiology across these diseases. CALY is associated with ADHD^34^, and 30-60% of tuberous sclerosis patients suffer from ADHD^35^. Additionally, calcyon haploinsufficiency has been proposed to be at least a partial cause of behavioral phenotypes identified in 10q26 deletion syndrome, as CALY is located at 10q26^36,37^. This suggests a role of insufficient or aberrant calcyon-mediated vesicle trafficking^38^ in TSC-associated ADHD. Relaxing the criteria of donor reproducibility but still maintaining reproducibility over batches admits a larger set of genes which are differentially expressed in eight or nine out of the ten possible donor pairs (**Figs. 5e, f**). Several ribosomal genes (RPL7, RPL10, RPL7A, RPS27, RPL28) are upregulated in this set. As mTOR regulates expression and translation of various ribosomal components, the dysregulation of ribosomal gene transcription is expected^39^, but further investigation is required to understand the role of these genes in TSC pathology.

## DISCUSSION

3D iPSC differentiation is sensitive to factors such as feeding regimen, media storage conditions, and liquid handling technique (**Supplementary Video 1**). Inspired by the contribution robotics has made to other aspects of drug discovery^40^, here we develop automated methods to ensure highly reproducible organoid culture (**Fig. 2**). Our platform, ORCA, also remedies a limitation of pooled culture protocols by enabling longitudinal tracking of individual organoids over time without compromising on culture scale. Comparing multiple assays on the same organoid, and at scale across organoids, permits detailed understanding of the sources of variability, and thus multi-donor, multi-clone, multi-batch phenotyping. Individual culture also enables tasks that would be impractical in pooled culture, such as efficient multifactorial protocol optimization (**Fig. 4)**.

The objective of phenotyping is to find disease signatures while factoring out biological (e.g., donor, clone) and technical (e.g., culture batch, assay variability) confounds (**Fig. 1b**). To address this, we developed the rank-to-group (RTG) score and are releasing an open-source Python package implementing it (https://github.com/System1Bio/rtg_score; **Fig. 3e-g)**. RTG scoring estimates the effect of any confounding variable for any assay. The only requirement is a distance metric between pairs of assay measurements. Since the method is agnostic to the confounds being analyzed and the assay, we anticipate it may be of broad utility in any attempt to characterize sources of variability in complex data sets.

Our work builds on previous studies characterizing organoid variability, such as the detailed analysis of developmental variation performed in kidney organoids by scRNA-seq^41^. Here, we have expanded such characterization to multiple assay modalities using the RTG score applied to a larger dataset spanning many donors, clones, and batches. The substantial contribution of batch, donor, clone, and their interactions to organoid differentiation suggests that phenotyping must be verified as robust across these confounds. Failure to do so opens two potential sources of error: first, attempts at measuring disease-associated signals may be overwhelmed by these confounds; and second, a proposed disease phenotype may spuriously arise from donor, clone, or batch variability. Importantly, we found that variability structure was matched across multiple assays (**Fig. 3e-g**) suggesting that automatable, non-destructive assays (e.g., brightfield imaging) can be used to track organoid variability just as well as destructive, resource-intensive assays (e.g., scRNA-seq).

TSC pathologies can present at 23 weeks gestation, motivating the urgency for prenatal therapy before permanent neurological deformities occur^42^. In recent work, disease phenotypes were only detected late in spheroid or organoid culture (>100 days)^19,20^, or under additional biallelic mosaic inactivation^20^. In one study, heterozygous TSC2^+/−^ mutations did not display detectable phenotypes^20^. By contrast, we detect a phenotype in heterozygous TSC^+/−^ organoids much earlier (day 35). Recent studies of iPSC-derived neural progenitor cells in 2D culture found TSC2^+/−^ mutants with developmental changes ^43^, but minimal differences between hESC-derived post-mitotic neurons in TSC^+/−^ and TSC^−/−^ mutants^44^. These 2D studies suggest less severe disease phenotypes may be present but hard to detect. Our phenotypes identified in heterozygous TSC2^+/−^ organoids (**Fig. 5**) suggests reevaluation of the hypothesis that a second-hit somatic mutation is required for the emergence of disease pathology and opens the door for early detection and treatment of the disease. Further research is warranted to explore the connection between the early disease phenotypes detected here, the developmental and morphological phenotypes identified later in development by Eichmuller et al.^19^, and those identified in 2D culture.

Successful *in vitro* disease modeling is a first and critical step for therapeutic development. Deep phenotyping can reveal more subtle and more novel signatures of disease that may be missed by more targeted approaches. Given the systems level nature of many neurodevelopmental diseases, the coordinated rescue of multiple phenotypes with a therapeutic intervention represents stronger evidence for advancing towards clinical development. Unbiased deep phenotypes identified in PDOs should also be of transformative value in later stages of drug discovery, including target identification, drug screening, drug optimization, biomarker development, toxicity testing, patient stratification, and clinical trial design. A scaled culture and assay system combined with large-scale data management and statistical analysis methods will be critical to enable all of these endeavors.

## METHODS

### Generation of iPSCs by fibroblast reprogramming

Fibroblasts were cultured in DMEM/F12 with 10% FBS and 1% NEAA, and passaged using TryPLE, and reprogrammed using Sendai virus (Thermo, Cytotune 2.0). Briefly, fibroblasts were plated at 7.5e3 cells / cm^2^ and 2.0e4 cells / cm^2^, and transduced the following day with recommended viral titers (KOS MOI = 5, hc-Myc MOI = 5, hKlf4 MOI = 3). Transduced cells were fed every other day with fibroblast medium until day 7. Cells were passaged using TryPLE on day 7, counted and replated at 2e3 cells / cm^2^ to 4e3 cells / cm^2^ in fibroblast medium. The following day, medium was replaced with Essential 8 Flex (Thermo) and fed daily until colonies appeared. Colonies were selected based on morphology and picked into Matrigel (Corning)-coated 24 well plates in StemFlex medium containing 10 uM ROCK inhibitor. Clones were then passaged 1:2 using EDTA until they reached 6 well formats (10 cm^2^), and banked into 3-4 vials using mFreSR (Stemcell Technologies). Clearance of Sendai virus was confirmed using qPCR (**Supplementary Table 4**). After Sendai clearance, iPSCs from each clone were expanded into multiple 10 cm dishes and banked into 20+ vials to make up a master bank. Details of all iPSC clones used are listed in **Supplementary Table 5**. TSC donor clinical information is listed in **Supplementary Table 2**.

### Individual well culture of organoids

Organoids were grown in individual wells using both unpatterned (UP) and directed, dorsal forebrain protocols (DFP). Briefly, our UP largely follows the protocol presented in Lancaster & Knoblich (2014)^22^ with the notable exception that we use individual wells as opposed to bioreactors. Master-banked control, TSC1^+/−^ and TSC2^+/−^ iPSCs were thawed concurrently, and passaged twice before seeding. Prior to seeding, iPSCs were dissociated when 50–80% confluent using Accutase (ThermoScientific) and low attachment U-bottom 96-well plates (Corning) at 1.0e4 cells / well. Healthy and TSC patient iPSCs were spatially intermixed on the same plate to account for any well, plate and downstream assay biases (e.g. temperature, humidity and imaging artifacts) (**Supplementary Fig. 4**). Organoids were embedded in 100% Matrigel Basement Membrane (Corning) on day 7. Following embedding, organoids were cultured in neural induction medium until day 13, when they were fed with differentiation medium. On day 21 they were transferred to 24-well plates and cultured until experimental endpoints, which could be beyond day 100. Protocol evolution experiments are readily enabled by the plate-based format and two DFPs are presented. Both DFP1 and DFP2 are based on neural induction via dual-SMAD inhibition as well as Wnt signaling inhibition for rostralization of the neuroepithelium. Full methods details are available in Supplementary Table 1.

### Automation-assisted Orgbook metadata tracking

For each individual organoid we culture, we comprehensively track the following categories of information across the workflow and data hierarchy:

1. Metadata about the patient donor including clinical history (known mutation sequences, age at disease onset, clinical symptoms and severity, treatment history, family relationships to other donors, and outcome measure) and sample acquisition information (age at biopsy, biopsy location, and primary cell type of biopsy)
2. External source of primary patient samples (e.g., fibroblast or blood sample) or iPSC clone (including catalog and lot numbers)
3. Reprogramming method, karyotyping, genomic integrity,Single Nucleotide Polymorphism (SNP) arrays, and metadata of any genetic modifications (e.g., gRNA, vector and plasmid used) of iPSC clones
4. Liquid nitrogen (LN2) inventory, iPSC master bank information, metadata on culture history including plate and well of each clone in culture, passage number, confluence at seeding, passage split ratios or cell counts and viability
5. Well, plate, and incubator locations of organoids as well as handling, transfer, harvests and dissociation events throughout their lifetime
6. Organoid culture protocol details including concentrations and timings of all media components and additives
7. Vendor, catalog number, and lot number of acquired compounds; compound treatment duration and concentration
8. Metadata associated with automated media sampling (driven by Hamilton Microlab STARlet via a customized hit-picking schema) and regular feeding events (driven by BioTek EL406 Washer Dispenser system with BioStack Microplate Stacker). All these routine liquid handling events were recorded by their unique run IDs that were tracked by Orgbook as well.
9. Daily or weekly automated imaging data including parameters and time of acquisition for every microscope used during the lifetime of the organoids (for both fluorescence and brightfield microscopy). Robotic arm (PA PF-400, PreciseAutomation), plate hotel, and custom-build, automated microscope system (ASI/Applied Scientific Instrumentation) were integrated to enable hands-free plate loading and communication with Orgbook (e.g., tracking the completion of daily imaging event for a specific barcoded organoid plate).
10. Metadata & sample prep QC associated with all assays (for example: for scRNA-seq: kit version, microfluidic lane, microfluidic chip, cell counts and viability during prep, BioAnalyzer library size quantification, KAPA qPCR library quantification; for immunofluorescence imaging: primary and secondary antibodies or chemical stains used, dilutions, lot numbers, freeze blocks, imaging parameters)

These data are easily accessible to all scientists via Orgbook, our relational database and internal web interface that supports experiment design, automation scheduling, data querying, and data visualization.

### qPCR

RNA was extracted from organoids using the Simply RNA Tissue kit (Promega, AS1340) on the Maxwell RSC 48 instrument (Promega). cDNA was synthesized using SuperScript III (Thermo, 18080051) with random hexamer primers. Amplification and quantitation was performed using PerfeCTa^®^ SYBR^®^ Green SuperMix (QuantaBio, 95056) on the QuantStudio 5 (Thermo). Primers used are shown in **Supplementary Table 4**.

### Flow cytometry

The organoids cultured from both 24w and 96w plates were dissociated into single cell suspension using a commercially available papain digestion kit (Worthington-biochem, LK003150). Live cells and dead cells from the dissociated organoids were stained with Calcium Green AM (ThermoFisher C3011MP) and Draq7 (Biolegend 424001), respectively, immediately before running flow cytometric analysis on Attune NxT Flow Cytometer (ThermoFisher). Analysis was performed using FlowJo (Becton Dickinson).

### Brightfield imaging

Brightfield images of organoids were obtained using either IN Cell Analyzer 2200, IN Cell Analyzer 2500, or EVOS FL Auto 2 at 4X magnification. Multiple images of each organoid were taken, either daily or weekly.

### Brightfield image analysis

Multiple z-planes are collapsed to a single image by weighted mixing of individual images so in each region an image with more high-frequency content is given higher weight. Conversion to grayscale is applied if necessary. Convolutional neural networks are used to detect and segment organoids in images (U-Net^45^) and to project cropped and resized images to embedding dimensions (ResNet-18^46^). Model embeddings are projected onto the sphere and ArcFace^47^ loss is minimized during training using Adam optimizer^48^. This forces embeddings of images from the same organoid, taken at different ages and augmented to be closer than embeddings from different organoids. Hard mining is used to prefer cases with the same organoid being imaged using different microscopes or in different plates to force the model to rely on biologically relevant features. Further details on the machine learning model are provided in the Supplementary Text.

QPCR values were predicted directly from embeddings by regularized linear OLS model using leave-organoid-out cross-validation.

### Analysis of contributing factors (RTG score)

The contribution of different confounding factors to qPCR gene expression values (**Fig. 3e**) using the RTG score is estimated as shown in **Supplementary Fig. 5.** Here we describe how the RTG score is computed for a given confounder (or set of confounders). Consider an arbitrary query organoid in the dataset. The RTG score for this organoid is defined as the probability that a random organoid with shared confounders (e.g., same donor) is more similar to the query organoid than a random organoid with different confounders (e.g., different donor). To compute this probability, we rank all other organoids according to their similarity to the query organoid (e.g., by Euclidean distance between their qPCR gene expression vectors), and then compute how well the similarity rank of each organoid maps to whether or not it shares the confounder(s) in question with the query organoid. This can be efficiently computed using an area under the ROC curve (AUROC). Additionally, exclusion criteria can be used to ensure the RTG score for a given confounder is not better explained by another confounder. For example, the ‘exclude-same-clone’ evaluation strategy used to evaluate the donor RTG score implies all images from the same clone as the query organoid are omitted from the similarity ranking. The final RTG score for a given confounder is the average RTG score computed over all query organoids.

Analysis of other modalities uses the same approach (**Figs. 3f, g**), but similarity between organoids is defined differently (Euclidean distance for brightfield image embeddings, and Hellinger distance between cell type distributions for scRNA-seq). The RTG score is agnostic to the choice of distance metric, so suitable distance metrics may be chosen that are appropriate for the data type in question. A complete Python implementation of the RTG score is provided here: https://github.com/System1Bio/rtg_score.

### Brightfield-based organoid size estimation

Size is computed using an automated segmentation model described above, that estimates the area the organoid occupies in the well from a brightfield image.

Organoid cross-sectional area was calculated from top-down brightfield images and used to fit organoid growth curves. Growth curves were estimated from a mixed-effects spline regression model (a generalized additive model (GAM) that uses a series of local polynomials to approximate arbitrary curves when summed together, and has been used to model growth dynamics^49^). We used natural cubic splines over the domain [0, 60] with knots as 20 and 40 days (effective DOF = 4).

For the mixed-effects portion of the model, we set organoids nested within their corresponding donors as random effects, and allowed the coefficients for the spline basis functions to vary. We chose this hierarchical structure as individual organoids were measured at multiple time points, and organoids derived from the same donor should not be considered independent from one another. Inference was performed using the lme4 R package^50^ via a python wrapper (Pymer4)^51^.

### Calcium imaging - experimental protocol

Prior to imaging, fresh calcium-sensitive dye (Fluo-4 direct, ThermoFisher, CAT # F10471) was prepared according to the recommended protocol. Organoids to be imaged were then transferred to 24-well imaging plates (Costar, Not-treated Polystyrene #3738) and washed twice with fresh media. Then, 400uL of media + dye was added to each well and allowed to incubate for 30 minutes within a standard cell culture incubator. After 30 minutes, organoids were placed into a dedicated, heated environment on a custom-built inverted epifluorescence microscope (ASI). Imaging was performed using a Lumencor Spectra X (Leica Microsystems) light source set to 15% power for 6 minutes at a frame rate of 10 Hz.

### Calcium imaging - video processing & analysis

Raw H5 videos were automatically processed by an ORCA software pipeline. Briefly, each video file underwent a series of custom preprocessing stages that (a) cropped out empty portions of the well, (b) globally aligned the frames to account for slow drift, (c) removed rigid and non-rigid motion of the organoid tissue, (d) eliminated floating debris, and (e) compensated for bleaching. Next, preprocessed videos were manually inspected and flagged if any of the preprocessing stages had failed (e.g. excessive motion, debris, etc.). Finally, putative neurons were identified from the preprocessed videos passing QC using CaImAn^52^, which were then refined using custom filters. Calcium dynamics and inferred spikes for the remaining neurons were stored in our database for offline analysis.

Traces and spikes for the identified neurons were downloaded and analyzed via custom Python code. Distributions for firing rates and inter-spike intervals were computed directly from the inferred spike times. For the PSD decay constants (**Fig. 2f**), first the power-spectral density was computed from the autocorrelation of each neuron’s calcium trace using Welch’s method, and then an exponential curve was fit to the computed PSD (on a semi-log axis) and the decay constant was stored.

### Organoid dissociation and library prep for scRNA-seq

Individual brain organoids were dissociated into a single-cell suspension using Accumax. Dissociated cells were resuspended in PBS+2% BSA, filtered through 40 μm cell strainers (Flowmi, H13680-0040), counted, and pooled across donors. Approximately 42,000 cells per channel (to give estimated recovery of 10,000 cells per channel) were loaded onto a Chromium Single Cell 3’ Chip and processed through the Chromium controller to generate single-cell gel beads in emulsion (GEMs). scRNA-seq libraries were prepared with the Chromium Single Cell 3’ Library & Gel Bead Kit v2 or v3. Libraries were pooled and sequenced on a NovaSeq 6000 (Illumina) with 26 bases for read 1, 98 bases for read 2 and 8 bases for index 1.

### scRNA-seq analysis

Reads were trimmed and aligned using CellRanger 3.0.2. In order to reconstruct donor identity for droplets and identify multiplets, a novel genotype-based demultiplexing algorithm was used. Initial information about donors’ genotypes was collected using bead arrays (Infinium Global Screening Array-24 Kit, v2 or v3) in the form of SNVs. Genotypes were later refined from scRNA-seq data by detecting new SNVs and correcting those provided by GSA using expectation-maximization approach. An algorithm is also set up to detect the presence of cells from donors not pooled to a library (used to detect contamination). Droplets which have posterior probability greater than 99% being assigned to a single donor, are analyzed for DEG.

Cells with over 15% mitochondrial reads, fewer than 300 expressed genes, or more than 10,000 expressed genes are filtered out. Seurat v3^53,54^ is used to identify the top 2000 variable genes and perform cell-wise dimensionality reduction using PCA on those 2000 genes. Cells are clustered into putative cell types using Louvain clustering (resolution=0.25, epsilon=0) with each cell represented by its top 15 PCs. Cell types are qualitatively defined using the top 10 over-expressed genes identified in each cluster via one-vs-rest differential expression (Wilcoxon rank-sum test) (**Supplementary Table 3**).

Two-dimensional scatter plots of cells are generated using UMAP on the top 30 PCs (n_neighbors=10, min_dist=0.5). Differentially expressed genes for disease phenotyping are identified by Wilcoxon rank-sum test.

We perform differential expression separately in each computationally-identified cluster (c_i_), in each batch (bj), between each pair of donors where one donor (D_WT_) is wild-type and the other donor (D_MUT_) has TSC. This results in a set of genes differentially expressed in particular clusters, between particular donors, in a particular batch: <gene, c_i_, bj, [D_WT_, D_MUT_]_k_> → <p_value, fold_change>. We reject any that have Bonferroni-adjusted *p*-value > 0.05 or absolute value of log_2_ fold change < 0.25. Next we reject any <gene, c_i_, b_j_, [D_WT_, D_MUT_]_k_> where it is significant in some batch b_i_, but not significant or significant in the other direction in another batch b_j_. This leaves a set of <gene, c_i_, [D_WT_, D_MUT_]_k_> that are consistent across all measured batches. We reject any <gene, c_i_> where <gene, c_i_, [D_WT_, D_MUT_]_k_> is significant in one direction for some donor pair k, but <gene, c_i_, [D_WT_, D_MUT_]_l_> is significant in the other direction for some other donor pair. This leaves a set of <gene, c_i_> that are consistent in both batches, in at least one donor pair, and not contradictorily differentially expressed in another donor pair. We plot these according to the number of donor pairs (out of a maximum of # of WT donors * # of MUT donors) in which <gene, c_i_> is statistically significant.

### Immunofluorescence staining and imaging

Organoids were transferred to 1.5ml tubes, rinsed with PBS, and fixed in 4% paraformaldehyde (Electron Microscopy Sciences, 15710) for 1 hour. After fixation, organoids were rinsed 3x with PBS, and incubated with 30% w/v sucrose solution overnight. The following day, samples were embedded into O.C.T. (Tissue Tek, 4583) and frozen at −80°C overnight. Embedded blocks were sectioned into 25 μm slices using a NX-70 Cryostar Cryostat (Thermo), and dried at room temperature overnight. Slides were stained on Sequenza racks (Thermo) the following day. Briefly, slides were rinsed with PBS, then incubated in 0.25% Triton X-100 with 4% Donkey serum (EMD Millipore, S30) for 1 hour, followed by incubation in primary antibody overnight. After rinsing 5x with PBS, slides were incubated in secondary antibodies for 2 hours at room temperature, stained with DAPI, and rinsed again 5x with PBS. Slides were then mounted, and imaged using Zeiss Axio Scan Z1 with 20x / 0.8 objective, and stitched together by the Zeiss Zen software. Post-image processing utilized FIJI (NIH) and Adobe Photoshop CC.

### Immunofluorescence phenotyping analysis

Images are then downsampled twice to reduce noise. ResNet-like architecture is employed with max-pooling replaced by striding in convolutions. Regions are picked randomly within a pool of images while ensuring no overlap. Figure 4B shows the training scheme: each region (large purple square) is associated with several clusters (e.g. C1, C2, C3) (3 clusters per region in our experiments). During training small tissue patches (small red squares) are converted by model into embeddings. Optimization target is set to attract embedding of each patch to one of clusters while repulsing from all clusters corresponding to other regions. Several clusters per region is a way to allow prediction of different tissue within a region and it resolves potential problems with boundaries between different tissue types.

Additional augmentations (affine transforms, brightness and contrast alterations) are applied to achieve model stability towards technical factors. Dataset contains organoid slices that were imaged at different timepoints or at different depths, and an additional term in the loss corresponds to distance between embeddings of the same patch images. To predict if a slice comes from a TSC organoid a regularized logistic regression was trained on the top of embeddings using leave-one-donor-out cross-validation scheme. Further details on the machine learning model are provided in the Supplementary Text.

## Author Contributions

GS Escola, S Kato, G Skibinski, K Shah, R Bedi, A Rogozhnikov, Z Tong, B Rash, J Sorokin, O Wueseke, and J Hilliard conceived of and designed experiments. G Skibinski, K Shah, B Rash, M Stanton, CY Lee, PK Hsu, O Wueseke, J Hilliard, S Drusinsky, D Quang, I Oumzil, J Bergamaschi, J Paek, C Apaydin, K Li, G Treiman, A Piepho, L Enriquez, P Taylor, R Blattner, and A Batarse performed laboratory experiments. A Rogozhnikov, R Bedi, J Sorokin, N Young, P Ramkumar, A McKay, D Grayson, and M Schultz built machine learning models and performed data analysis. A Lash, A Bosshardt, C Johnson, S Brown, and D Flanzer built OrgBook, laboratory automation software, and data pipelines. Z Tong, J Nicola, C Castrillo, E Mount, S Poust, and D Flanzer built the ORCA automation infrastructure. R Bedi, P Ramkumar, A Rogozhnikov, K Shah, GS Escola, and S Kato wrote the manuscript; with input from G Skibinski, M Stanton, J Sorokin, B Rash, and Z Tong. All authors discussed results / reviewed the manuscript. GS Escola, S Kato, G Skibinski, D Flanzer, and D Chao oversaw the project.

## References

1. Yoon, S.-J. et al. Reliability of human cortical organoid generation. Nat. Methods 16, 75–78 (2019).

2. Lancaster, M. A. et al. Cerebral organoids model human brain development and microcephaly. Nature 501, 373–379 (2013).

3. Velasco, S. et al. Individual brain organoids reproducibly form cell diversity of the human cerebral cortex. Nature 570, 523–527 (2019).

4. Bershteyn, M. et al. Human iPSC-Derived Cerebral Organoids Model Cellular Features of Lissencephaly and Reveal Prolonged Mitosis of Outer Radial Glia. Cell Stem Cell 20, 435–449.e4 (2017).

5. Kim, J., Koo, B.-K. & Knoblich, J. A. Human organoids: model systems for human biology and medicine. Nat. Rev. Mol. Cell Biol. (2020) doi:10.1038/s41580-020-0259-3.

6. Trujillo, C. A. et al. Complex Oscillatory Waves Emerging from Cortical Organoids Model Early Human Brain Network Development. Cell Stem Cell 25, 558–569.e7 (2019).

7. Mansour, A. A. et al. An in vivo model of functional and vascularized human brain organoids. Nat. Biotechnol. 36, 432–441 (2018).

8. Pacitti, D., Privolizzi, R. & Bax, B. E. Organs to Cells and Cells to Organoids: The Evolution of Central Nervous System Modelling. Front. Cell. Neurosci. 13, 129 (2019).

9. Fang, Y. & Eglen, R. M. Three-Dimensional Cell Cultures in Drug Discovery and Development. SLAS Discov 22, 456–472 (2017).

10. Volpato, V. & Webber, C. Addressing variability in iPSC-derived models of human disease: guidelines to promote reproducibility. Dis. Model. Mech. 13, (2020).

11. Quadrato, G. et al. Cell diversity and network dynamics in photosensitive human brain organoids. Nature 545, 48–53 (2017).

12. Cheadle, J. P., Reeve, M. P., Sampson, J. R. & Kwiatkowski, D. J. Molecular genetic advances in tuberous sclerosis. Hum. Genet. 107, 97–114 (2000).

13. Thiele, E. A. Managing and understanding epilepsy in tuberous sclerosis complex. Epilepsia 51 Suppl 1, 90–91 (2010).

14. Jansen, F. E. et al. Cognitive impairment in tuberous sclerosis complex is a multifactorial condition. Neurology 70, 916–923 (2008).

15. Crino, P. B., Nathanson, K. L. & Henske, E. P. The tuberous sclerosis complex. N. Engl. J. Med. 355, 1345–1356 (2006).

16. Kwiatkowski, D. J. Animal models of lymphangioleiomyomatosis (LAM) and tuberous sclerosis complex (TSC). Lymphat. Res. Biol. 8, 51–57 (2010).

17. Blair, J. D. & Bateup, H. S. New frontiers in modeling tuberous sclerosis with human stem cell-derived neurons and brain organoids. Dev. Dyn. 249, 46–55 (2020).

18. Mizuguchi, M. & Takashima, S. Neuropathology of tuberous sclerosis. Brain Dev. 23, 508–515 (2001).

19. Eichmüller, O. L. et al. Cerebral organoid model reveals excessive proliferation of human caudal late interneuron progenitors in Tuberous Sclerosis Complex. Neuroscience (2020).

20. Blair, J. D., Hockemeyer, D. & Bateup, H. S. Genetically engineered human cortical spheroid models of tuberous sclerosis. Nat. Med. 24, 1568–1578 (2018).

21. Martin, K. R. et al. The genomic landscape of tuberous sclerosis complex. Nat. Commun. 8, 15816 (2017).

22. Lancaster, M. A. & Knoblich, J. A. Generation of cerebral organoids from human pluripotent stem cells. Nat. Protoc. 9, 2329–2340 (2014).

23. Camp, J. G. et al. Human cerebral organoids recapitulate gene expression programs of fetal neocortex development. Proc. Natl. Acad. Sci. U. S. A. 112, 15672–15677 (2015).

24. Nordström, U., Jessell, T. M. & Edlund, T. Progressive induction of caudal neural character by graded Wnt signaling. Nat. Neurosci. 5, 525–532 (2002).

25. Kiecker, C. & Niehrs, C. A morphogen gradient of Wnt/beta-catenin signalling regulates anteroposterior neural patterning in Xenopus. Development 128, 4189–4201 (2001).

26. Jerber, J. et al. Population-scale single-cell RNA-seq profiling across dopaminergic neuron differentiation. bioRxiv 2020.05.21.103820 (2020) doi:10.1101/2020.05.21.103820.

27. Amin, N. D. & Paşca, S. P. Building Models of Brain Disorders with Three-Dimensional Organoids. Neuron 100, 389–405 (2018).

28. Kadoshima, T. et al. Self-organization of axial polarity, inside-out layer pattern, and species-specific progenitor dynamics in human ES cell–derived neocortex. Proc. Natl. Acad. Sci. U. S. A. 110, 20284–20289 (2013).

29. Mariani, J. et al. Modeling human cortical development in vitro using induced pluripotent stem cells. Proc. Natl. Acad. Sci. U. S. A. 109, 12770–12775 (2012).

30. Hanashima, C., Shen, L., Li, S. C. & Lai, E. Brain Factor-1 Controls the Proliferation and Differentiation of Neocortical Progenitor Cells through Independent Mechanisms. J. Neurosci. 22, 6526–6536 (2002).

31. Pham, X. et al. The DPYSL2 gene connects mTOR and schizophrenia. Transl. Psychiatry 6, e933 (2016).

32. Liu, Y. et al. Functional variants in DPYSL2 sequence increase risk of schizophrenia and suggest a link to mTOR signaling. G3 5, 61–72 (2014).

33. Takata, A. et al. Integrative Analyses of De Novo Mutations Provide Deeper Biological Insights into Autism Spectrum Disorder. Cell Rep. 22, 734–747 (2018).

34. Laurin, N. et al. Association of the calcyon gene (DRD1IP) with attention deficit/hyperactivity disorder. Mol. Psychiatry 10, 1117–1125 (2005).

35. D'Agati, E., Moavero, R., Cerminara, C. & Curatolo, P. Attention-deficit hyperactivity disorder (ADHD) and tuberous sclerosis complex. J. Child Neurol. 24, 1282–1287 (2009).

36. Lin, S. et al. Chromosome 10q26 deletion syndrome: Two new cases and a review of the literature. Mol. Med. Rep. 14, 5134–5140 (2016).

37. Distal 10q monosomy: New evidence for a neurobehavioral condition? Eur. J. Med. Genet. 57, 47–53 (2014).

38. Shi, L., Muthusamy, N., Smith, D. & Bergson, C. Dynein binds and stimulates axonal motility of the endosome adaptor and NEEP21 family member, calcyon. Int. J. Biochem. Cell Biol. 90, 93–102 (2017).

39. Mayer, C. & Grummt, I. Ribosome biogenesis and cell growth: mTOR coordinates transcription by all three classes of nuclear RNA polymerases. Oncogene 25, 6384–6391 (2006).

40. Schneider, G. Automating drug discovery. Nat. Rev. Drug Discov. 17, 97–113 (2018).

41. Phipson, B. et al. Evaluation of variability in human kidney organoids. Nat. Methods 16, 79–87 (2019).

42. Prabowo, A. S. et al. Fetal brain lesions in tuberous sclerosis complex: TORC1 activation and inflammation. Brain Pathol. 23, 45–59 (2013).

43. Zucco, A. J. et al. Neural progenitors derived from Tuberous Sclerosis Complex patients exhibit attenuated PI3K/AKT signaling and delayed neuronal differentiation. Mol. Cell. Neurosci. 92, 149–163 (2018).

44. Costa, V. et al. mTORC1 Inhibition Corrects Neurodevelopmental and Synaptic Alterations in a Human Stem Cell Model of Tuberous Sclerosis. Cell Rep. 15, 86–95 (2016).

45. Ronneberger, O., Fischer, P. & Brox, T. U-Net: Convolutional Networks for Biomedical Image Segmentation. arXiv [cs.CV] (2015).

46. He, K., Zhang, X., Ren, S. & Sun, J. Deep Residual Learning for Image Recognition. arXiv [cs.CV] (2015).

47. Deng, J., Guo, J., Xue, N. & Zafeiriou, S. ArcFace: Additive Angular Margin Loss for Deep Face Recognition. arXiv [cs.CV] (2018).

48. Kingma, D. P. & Ba, J. Adam: A Method for Stochastic Optimization. arXiv[cs.LG] (2014).

49. Grajeda, L. M. et al. Modelling subject-specific childhood growth using linear mixed-effect models with cubic regression splines. Emerg. Themes Epidemiol. 13, 1 (2016).

50. Bates, D., Mächler, M., Bolker, B. & Walker, S. Fitting Linear Mixed-Effects Models Using lme4. Journal of Statistical Software, Articles 67, 1–48 (2015).

51. Jolly, E. & Rose, S. A. ejolly/pymer4: JOSS Complete. (2018). doi:10.5281/zenodo.1523205.

52. Giovannucci, A. et al. CaImAn an open source tool for scalable calcium imaging data analysis. Elife 8, (2019).

53. Stuart, T. et al. Comprehensive Integration of Single-Cell Data. Cell 177, 1888–1902.e21 (2019).

54. Butler, A., Hoffman, P., Smibert, P., Papalexi, E. & Satija, R. Integrating single-cell transcriptomic data across different conditions, technologies, and species. Nat. Biotechnol. 36, 411–420 (2018).

