## Supplemental figures and methods for "Optimization and scaling of patient-derived brain organoids uncovers deep phenotypes of disease"

### Supplementary Information

#### SUPPLEMENTARY VIDEOS

##### *Supplementary Video 1* – Organoid feeding speeds

#### SUPPLEMENTARY FIGURES

##### *Supplementary Figure 1*

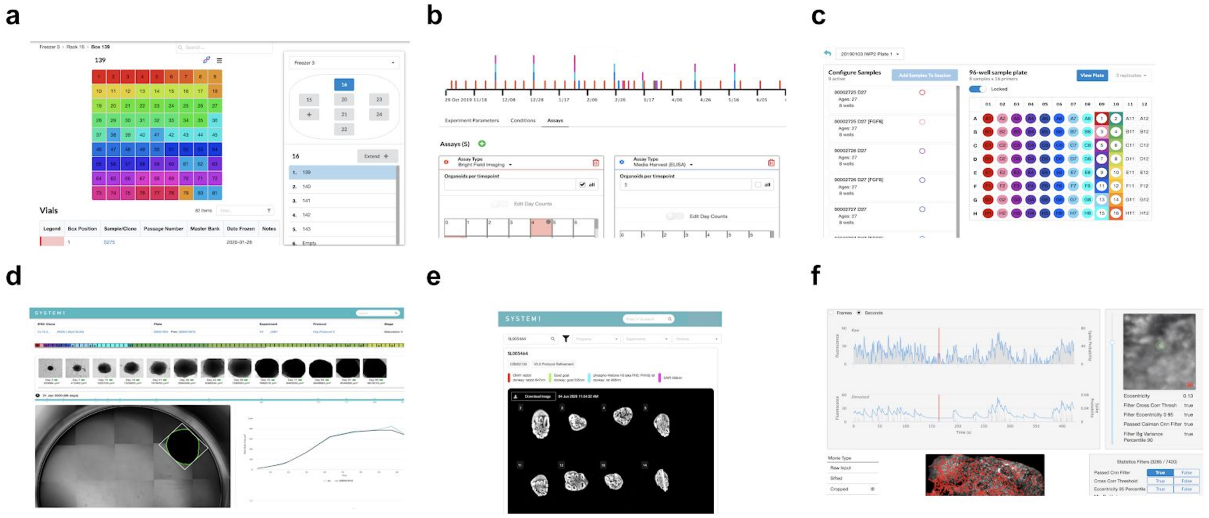

**Supplementary Figure 1** | Orgbook screenshots illustrating web applications for (a) biorepository management (b) experiment planning (c) automation control (d) experiment monitoring (e) data access and visualization and (f) data annotation and reporting.

##### *Supplementary Figure 2*

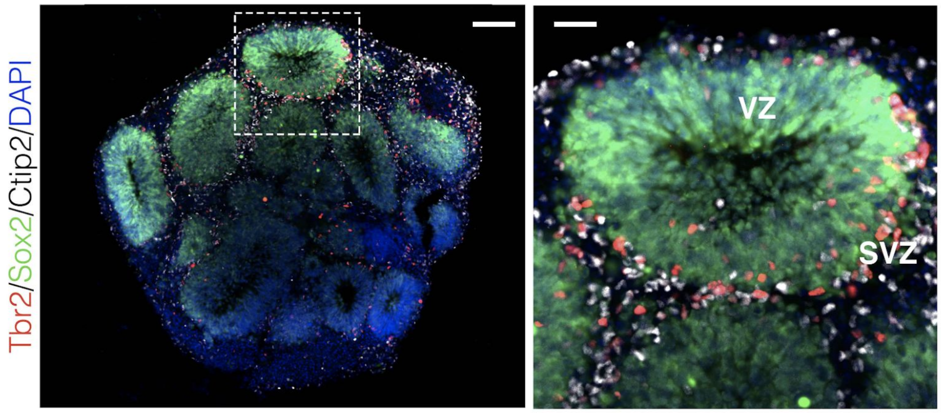

**Supplementary Figure 2** | (left) Immunofluorescence image of day 31 DFP organoid; (right) zoomed-in version of a bud from (left) with clear emergence of ventricular and subventricular zones (VZ, SVZ); scale bar is 100 μm (left), 30 μm (right)

#### *Supplementary Figure 3*

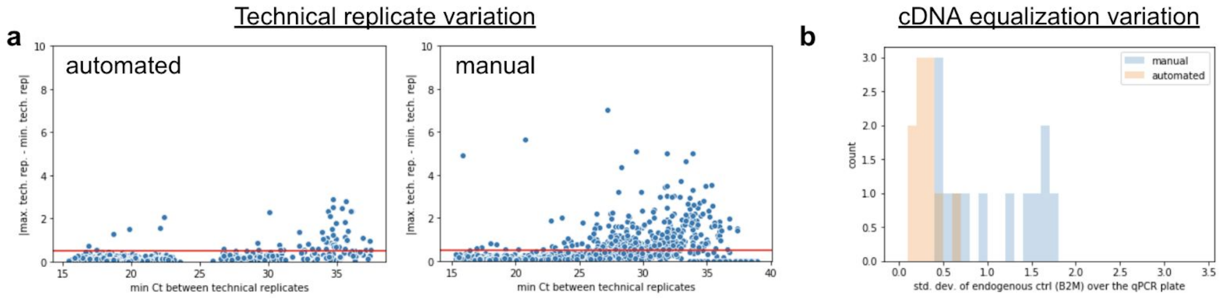

**Supplementary Figure 3** | **(a)** Maximum absolute difference between technical replicates as a function of minimum Ct across the same set of replicates for automated (left) and manual (right) implementations of qPCR. Each dot represents an organoid sample run across multiple replicates. Red line represents a maximum difference between any technical replicates of 1.0. (**b**) Distribution of Ct standard deviation across 12 qPCR 384-well plates for an endogenous control gene (B2M) for manual vs automated workflows.

#### *Supplementary Figure 4*

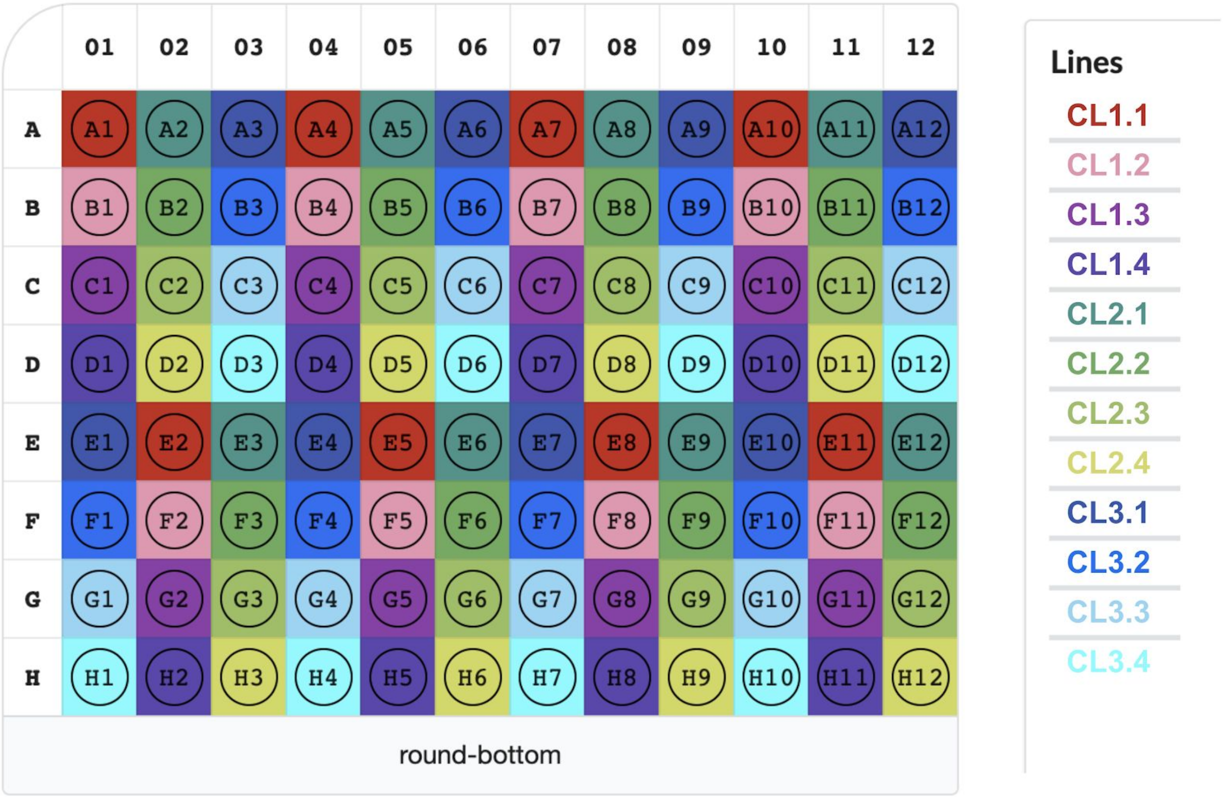

**Supplementary Figure 4** | An illustrative screenshot of Orgbook illustrating organoids from different donors and clones on the same plate as part of a single seeding batch. Here, n=12 unique iPSC clones from 3 donors were dissociated separately, and seeded in alternating columns of round-bottom 96 well plates. Eight organoids were seeded per clone per plate, as demonstrated above. This design accounts for intra-plate effects, as column-wise aspiration/feeding volume discrepancies would not be biased by clone. Healthy and disease patient-derived organoids (PDOs) are seeded on the same plate to ensure disease phenotypes are not biased by inter-plate effects including plate handling, environmental exposure, reagent lots and media preparations.

#### *Supplementary Figure 5*

Example of estimating RTG score of *donor* in the *exclude-same-clone* condition on this 6 organoid dataset:

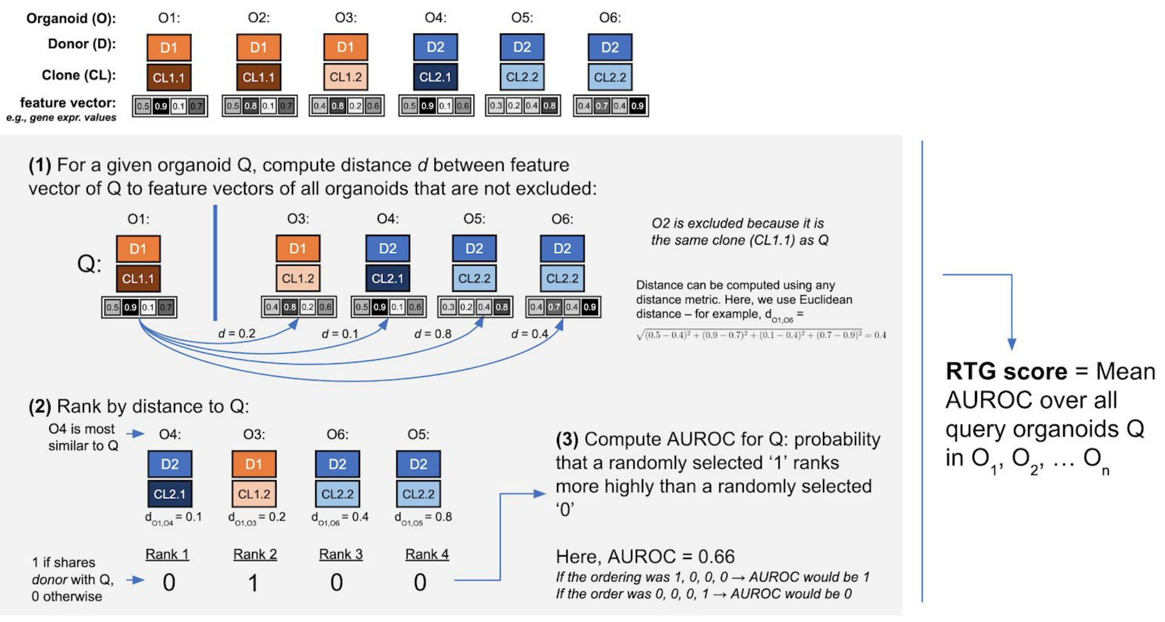

**Supplementary Figure 5** | An example of how the RTG score for the contribution of donor (using exclude-same-clone evaluation) to qPCR gene expression values is computed is provided here. Consider an arbitrary query organoid in the dataset (Q). First, organoids of the same clone as Q are excluded. This means that the final RTG score will not be influenced by organoids of the same clone meaning any reported contribution of donor will not actually reflect contribution of clone. Next, we compute the distance between Q and each non-excluded organoid. Here, this is shown using Euclidean distance between gene expression vectors, but in practice, could be any reasonable distance metric between any observed features. Next, we rank each organoid by its similarity to Q. Rank 1 is most similar. We compute the AUROC score that indicates how well the similarity rankings map to whether or not the donor property is shared.  The final RTG score for a given confounder is the average RTG score computed over all query organoids.

#### *Supplementary Figure 6*

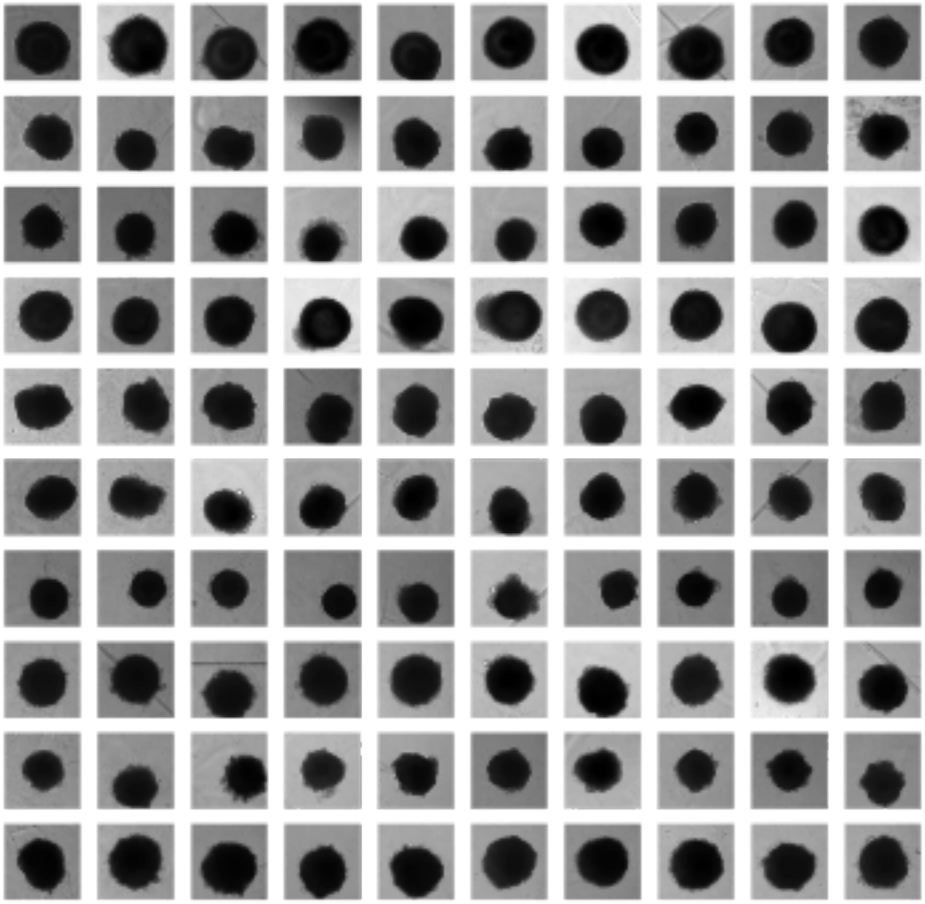

**Supplementary Figure 6** | Grid of DFP2 organoid images at day 14 from a single batch. Each row is a distinct clone.

#### *Supplementary Figure 7*

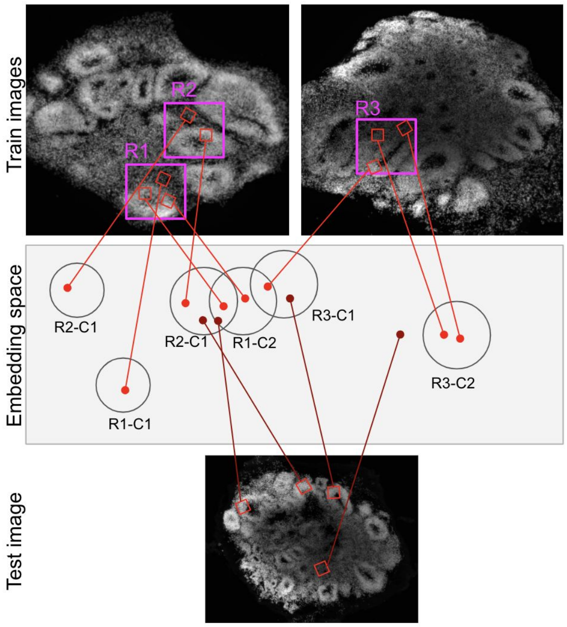

**Supplementary Figure 7** | Schematic of the self-supervised machine learning system used to train the feature extractor to learn vector embeddings of patches of DAPI-stained tissue by recognizing which point in embedding space (red) best represents a patch (red) in relation to other patches. Patches from three larger 225 μm x 225 μm regions (purple bounding boxes) are illustrated along with their corresponding points in embedding space.

Supplementary Table 1. Organoid protocols

| Inhibitors and morphogens | |
|---|---|
| **Compound** | **Range Tested** |
| RI Y27632 (1254 R&D Systems) | 5-80uM, D0-D5 |
| SB431542 (3014193 Biogems) | 2-10uM, D0-D7 |
| LDN193189 (1066208 Biogems) | 50nM-3.2uM, D0-D7 |
| IWP2 (6866167 Biogems) | 1-5uM, D0-D17 |
| FGF8b (100-25 Peprotech) | 200ng/mL, D5-7 |
| Cyc (Cyclopamine A; 4445185 Peprotech) | 1-20uM D5-17 |

| Timeline | UP |
|---|---|
| **Seeding (D0)** | 80% NI+hep, 20% StemFlex (SF) |
| **D2 Feed** | NI+hep |
| **D5 Feed** | NI+hep |
| **D7 Feed** | NI+hep, 100% matrigel embed |
| **D8-11 Feed** | NI+hep |
| **D12 Feed** | NI+hep |
| **D13-20 Feed** | Diff-A |
| **D21 Transfer Feed** | Diff-A |
| **D22-D41 Feed** | Diff-A |
| **D42-D50 Feed** | Diff+A |
| **D51+** | BrainPhys |

| Timeline | DFP1 | DFP2 |
|---|---|---|
| **Seeding (D0)** | 80%NI-hep, 20% SF, 40uM RI, 10uM SB, 100nM LDN, 2.5uM IWP2 | 80%NI-hep, 20% SF, 10uM RI, 10uM SB, 100nM LDN |
| **D2 Feed** | 100%NI-hep, 40uM RI, 10uM SB, 100nM LDN, 2.5uM IWP2 | 100%NI-hep, 10uM SB, 100nM LDN, 2.5uM IWP2 |
| **D5 Feed** | 100%NI-hep, 10uM SB, 100nM LDN, 2.5uM IWP2, 1uM Cyc | 100%NI-hep, 10uM SB, 100nM LDN, 2.5uM IWP2, 200ng/mL FGF8b |
| **D7 Feed** | NI-hep + 1uM Cyc | NI-hep, 100% matrigel embed |
| **D8-11 Feed** | NI-hep + 1uM Cyc | NI-hep |
| **D12 Feed** | NI-hep | NI-hep |
| **D13-20 Feed** | Diff-A | Diff-A |
| **D21 Transfer Feed** | Diff-A | Diff-A |
| **D22-29 Feed** | Diff-A | Diff-A |
| **D30-D38 Feed** | Diff-A + 1% GFR MG | Diff-A |
| **D39-D60 Feed** | Diff+A + 1% GFR MG | Diff+A |
| **D61-74 Feed** | Diff+A | Diff+A |
| **D75+** | BrainPhys | BrainPhys |

| Media Compositions | | |
|---|---|---|
| **NI (+ or - heparin (hep))** | **Diff (+ or - Vitamin A)** | **BrainPhys** |
| 1X DMEM/F12 | 0.5X DMEM/F12 | 1X BrainPhys |
| 1X N2 | 0.5X Neurobasal | 1X B27 + Vitamin A |
| 1X Glutamax | 0.5X N2 | 20ng/mL BDNF |
| 1X MEM-NEAA | 1X B27 (+ or - Vitamin A) | 20ng/mL GNDF |
| 1X Pen/Strep | 2.5ug/mL human insulin | 1X Glutamax |
| 1ug/ml Heparin | 1X Glutamax | 1X Pen/Strep |
| | 0.5X MEM-NEAA | |
| | 12.5mM HEPES (Diff+A only) | |
| | 1X Pen/Strep | |

| cell_line_donor_id | cell_line_sample_catalog_number | mutated_gene | cell_line_donor_disease_genetic_mutation | clinical_notes |
|---|---|---|---|---|
| 12 | GM03958 | TSC2 | c.1564del (p.H522Tfs) | Clinically affected; adenoma sebaceum; ungual fibromas; shagreen patches; seizures; mental retardation; intracerebral calcifications; |
| 13 | GM04520 | TSC2 | Large deletion exon 1-14 | Clinically affected; seizures since age 4; mild mental retardation; adenoma sebaceum; hypopigmented macules on right flank and right distal thigh; cerebral calcifications on EMI scan; |
| 14 | GM06100 | TSC2 | c.2743-1G>A (IVS25-1G>A) | Clinically affected; seizures; hypomelanotic macules on left cheek; facial angiofibromas; confetti like hypopigmented macules scattered over lower legs; shagreen patches on posterior right shoulder, mid-upper back, mid-lower back, and right posterior thigh; adenoma sebaceum; two small scalp papules; ungual fibromas on left thumb and index finger and right thumb, index finger and middle finger; decreased visual acuity in right eye; |
| 15 | GM06102 | TSC2 | c.5228G>A (R1743Q) | Clinically affected; diagnosed at age 2; mental retardation with IQ of 66; adenoma sebaceum over the nose and cheeks in a butterfly distribution; seizures that began about 6 months of age; numerous, variable sized areas of depigmentation over the legs, arms and back; shagreen patch in midline of upper back; forehead plaque; abnormal EEG at age 7, which showed right frontotemporal and left occipital focal abnormalities; |
| 17 | GM06149 | TSC1 | c.2249G>A (W750X) | Clinically affected; seizures; hypopigmented macules on right shoulder, right thigh, left chest, right knee, lower back and right upper arm; cafe-au-lait spot on right wrist; confetti hypopigmentation over back and legs; myopia; at age 17 there were no shagreen patches, ungual fibromas or adenoma sebaceum; calcifications in brain; |

| Fig 2B | Cluster: | 0 | 1 | 2 | 3 | 4 | 5 | 6 | 7 |
|---|---|---|---|---|---|---|---|---|---|
| Top 10 genes per cluster (rank ordered) | 1 | SST | NRGN | HIST1H4C | FABP7 | SPARCL1 | NEUROD6 | TOP2A | TTR |
| | 2 | DLX6-AS1 | NNAT | HMGB2 | HES5 | COL3A1 | NEUROD2 | UBE2C | IGFBP7 |
| | 3 | PLS3 | TAC1 | TUBA1B | SFRP1 | PTN | LMO3 | CENPF | HTR2C |
| | 4 | CALB2 | MEIS2 | KIAA0101 | HOPX | ID3 | CSRP2 | PTTG1 | PCP4 |
| | 5 | ERBB4 | SIX3 | TYMS | VIM | APOE | NFIB | HMGB2 | CA2 |
| | 6 | SCGN | ZFHX3 | HMGN2 | SLC1A3 | COL1A1 | TBR1 | CCNB1 | CXCL14 |
| | 7 | MAF | PEG10 | DLX2 | PTN | VIM | GPM6A | NUSAP1 | SERPINF1 |
| | 8 | ARX | ZFHX4 | HES6 | PTPRZ1 | S100B | ZBTB18 | CKS2 | TRPM3 |
| | 9 | SYT1 | GAP43 | GAD2 | TTYH1 | METRN | NELL2 | CDK1 | RBP1 |
| | 10 | TCF4 | HMP19 | DUT | FAM181B | SPARC | SLA | CCNB2 | CCK |

| Fig 5F | Cluster: | 0 | 1 | 2 | 3 | 4 | 5 | 6 | 7 | 8 | 9 | 10 | 11 | 12 |
|---|---|---|---|---|---|---|---|---|---|---|---|---|---|---|
| Top 10 genes per cluster (rank ordered) | 1 | GNRH1 | NEUROD6 | TTR | C1orf61 | LHX1 | S100B | CENPF | COL1A2 | HES5 | HIST1H4C | S100A10 | NTRK2 | PRPH |
| | 2 | DLX6-AS1 | GPM6A | TPBG | FABP7 | STMN2 | LMO4 | UBE2C | COL3A1 | DLK1 | KIAA0101 | S100A6 | PTN | PPP1R17 |
| | 3 | DLX5 | EMX2 | WLS | SFRP1 | LHX9 | CAV1 | TOP2A | LGALS1 | FGFBP3 | TYMS | S100B | FGFBP3 | RGS10 |
| | 4 | PEG10 | TBR1 | RSPO3 | ID4 | ROBO3 | BCAN | PTTG1 | COL1A1 | RP3-395M20.12 | HIST1H1C | POSTN | ID2 | C11orf96 |
| | 5 | STMN2 | NTS | IGFBP5 | EMX2 | NEFM | SCRG1 | HMGB2 | LUM | NKX2-1 | HIST1H1B | OLFML2A | TTYH1 | ISL1 |
| | 6 | DCX | NRN1 | WNT4 | PTPRZ1 | TAGLN3 | NR2F2 | CCNB1 | MGP | SFTA3 | TOP2A | LGALS1 | NR2F1 | CADM1 |
| | 7 | SOX4 | NRXN1 | CLU | MASP1 | THSD7A | S100A10 | NUSAP1 | PRRX1 | LINC01551 | HMGN2 | S100A11 | RFX4 | NEUROD1 |
| | 8 | MEIS2 | NNAT | ID1 | TTYH1 | SYT4 | MPZ | CKS2 | TWIST1 | MGST1 | DEK | SPARC | GPM6B | PPP1R14A |
| | 9 | SYT1 | RTN1 | SLC2A1 | HES1 | SOX4 | SOX10 | KPNA2 | DCN | SMS | HIST1H1D | CDH6 | HES4 | C14orf132 |
| | 10 | DLX2 | KIF5C | RSPO2 | SOX2 | GAP43 | NPR3 | CCNB2 | VCAN | SOX2 | TUBA1B | BCHE | CYP26A1 | TLX3 |

| gene | forward_primer_sequence | reverse_primer_sequence |
|---|---|---|
| FOXG1 | tggcccatgtcgcccttcct | gccgacgtggtgccgttgta |
| EN1 | cgtggcttactccccattta | tctcgctgtctctccctctc |
| Pax6 | ttgctggcctgtcttctctg | tgaggccctggagaaagagt |
| WNT1 | cgatggtggggtattgtgaac | ccggattttggcgtatcagac |
| IRX3 | gagggaaacgcttatgggagc | cgccgtctaagttctccaaatc |
| GAPDH | ttgaggtcaatgaaggggtc | gaaggtgaaggtcggagtca |
| LMX1A | cgcatcgtttcttctcctct | cagacagacttggggctcac |
| TBP | gggcaccactccactgtatc | cgaagtgcaatggtctttagg |
| SeV-ALL | tggctaagaacatcggaagg | gttttgcaaccaagcactca |
| SeV-Oct3/4 | gaagcctttccccctgtctc | atcgaaggtgctcaacaaccc |
| SeV-Sox2 | cccagcactaccagagcg | tcgaaggtgctcaacaaccc |
| SeV-Klf4 | tgcgaccgagcattttccag | atcgaaggtgctcaacaaccc |
| SeV-c-Myc | tgcggaaacgacgagaacag | gaagggttttgggaggactcc |

| cell_line_donor_id | cell_line_clone_short_name | parent_sample_provider_name | reprogramming_provider_name | cell_line_sample_catalog_number | reprogramming_method |
|---|---|---|---|---|---|
| 3 | CL3.1 | Coriell | in-house | GM11270 | sendai-virus |
| 3 | CL3.2 | Coriell | in-house | GM11270 | sendai-virus |
| 3 | CL3.3 | Coriell | in-house | GM11270 | sendai-virus |
| 3 | CL3.4 | Coriell | in-house | GM11270 | sendai-virus |
| 3 | CL3.5 | Coriell | in-house | GM11270 | sendai-virus |
| 3 | CL3.6 | Coriell | in-house | GM11270 | sendai-virus |
| 6 | CL6.1 | Coriell | in-house | GM11273 | sendai-virus |
| 6 | CL6.2 | Coriell | in-house | GM11273 | sendai-virus |
| 6 | CL6.3 | Coriell | in-house | GM11273 | sendai-virus |
| 6 | CL6.4 | Coriell | in-house | GM11273 | sendai-virus |
| 6 | CL6.5 | Coriell | in-house | GM11273 | sendai-virus |
| 6 | CL6.6 | Coriell | in-house | GM11273 | sendai-virus |
| 8 | CL8.1 | Coriell | in-house | GM17567 | sendai-virus |
| 8 | CL8.2 | Coriell | in-house | GM17567 | sendai-virus |
| 8 | CL8.3 | Coriell | in-house | GM17567 | sendai-virus |
| 8 | CL8.4 | Coriell | in-house | GM17567 | sendai-virus |
| 8 | CL8.5 | Coriell | in-house | GM17567 | sendai-virus |
| 9 | CL9.1 | Coriell | in-house | GM17880 | sendai-virus |
| 9 | CL9.2 | Coriell | in-house | GM17880 | sendai-virus |
| 9 | CL9.3 | Coriell | in-house | GM17880 | sendai-virus |
| 9 | CL9.4 | Coriell | in-house | GM17880 | sendai-virus |
| 9 | CL9.5 | Coriell | in-house | GM17880 | sendai-virus |
| 9 | CL9.6 | Coriell | in-house | GM17880 | sendai-virus |
| 10 | CL10.1 | Coriell | in-house | GM25456 | sendai-virus |
| 10 | CL10.2 | Coriell | in-house | GM25456 | sendai-virus |
| 10 | CL10.3 | Coriell | in-house | GM25456 | sendai-virus |
| 10 | CL10.4 | Coriell | in-house | GM25456 | sendai-virus |
| 10 | CL10.5 | Coriell | in-house | GM25456 | sendai-virus |
| 10 | CL10.6 | Coriell | in-house | GM25456 | sendai-virus |
| 11 | CL11.1 | Coriell | in-house | GM21921 | sendai-virus |
| 11 | CL11.2 | Coriell | in-house | GM21921 | sendai-virus |
| 11 | CL11.3 | Coriell | in-house | GM21921 | sendai-virus |
| 12 | CL12.1 | Coriell | in-house | GM03958 | sendai-virus |
| 12 | CL12.2 | Coriell | in-house | GM03958 | sendai-virus |

| | | | | | |
|---|---|---|---|---|---|
| 12 | CL12.3 | Coriell | in-house | GM03958 | sendai-virus |
| 13 | CL13.1 | Coriell | in-house | GM04520 | sendai-virus |
| 13 | CL13.2 | Coriell | in-house | GM04520 | sendai-virus |
| 13 | CL13.3 | Coriell | in-house | GM04520 | sendai-virus |
| 14 | CL14.1 | Coriell | in-house | GM06100 | sendai-virus |
| 14 | CL14.2 | Coriell | in-house | GM06100 | sendai-virus |
| 14 | CL14.3 | Coriell | in-house | GM06100 | sendai-virus |
| 15 | CL15.1 | Coriell | in-house | GM06102 | sendai-virus |
| 15 | CL15.2 | Coriell | in-house | GM06102 | sendai-virus |
| 15 | CL15.3 | Coriell | in-house | GM06102 | sendai-virus |
| 16 | CL16.1 | Coriell | in-house | GM06121 | sendai-virus |
| 16 | CL16.2 | Coriell | in-house | GM06121 | sendai-virus |
| 16 | CL16.3 | Coriell | in-house | GM06121 | sendai-virus |
| 17 | CL17.1 | Coriell | in-house | GM06149 | sendai-virus |
| 17 | CL17.2 | Coriell | in-house | GM06149 | sendai-virus |
| 17 | CL17.3 | Coriell | in-house | GM06149 | sendai-virus |
| 22 | CL22.2 | Thermo Fisher Scientific | Thermo Fisher Scientific | A18944 | episomal |
| 25 | CL25.1 | Applied StemCell | Applied StemCell | ASE-9502 | episomal |
| 76 | CL76.1 | Coriell | Coriell | AG25370 | sendai-virus |
| 79 | CL79.1 | NINDS | NINDS | NDS00143 | episomal |
| 80 | CL80.1 | NINDS | NINDS | NDS00201 | episomal |
| 81 | CL81.1 | NINDS | NINDS | NDS00202 | episomal |
| 82 | CL82.1 | NINDS | NINDS | NDS00163 | episomal |
| 87 | CL87.1 | CIRM | CIRM | CW60124 | episomal |
| 88 | CL88.1 | CIRM | CIRM | CW20128 GG1 | episomal |
| 89 | CL89.1 | CIRM | CIRM | CW20144 CC1 | episomal |
| 90 | CL90.1 | CIRM | CIRM | CW20320 CC1 | episomal |
| 93 | CL93.1 | CIRM | CIRM | CW20051 CC1 | episomal |
| 95 | CL95.1 | CIRM | CIRM | CW20109 FF1 | episomal |
| 98 | CL98.1 | CIRM | CIRM | CW20106 FF1 | episomal |
| 101 | CL101.1 | CIRM | CIRM | CW60220 FF1 | episomal |
| 102 | CL102.1 | CIRM | CIRM | CW60428 CC1 | episomal |
| 103 | CL103.1 | CIRM | CIRM | CW10173 EE1 | episomal |
| 110 | CL110.2 | Rett Syndrome Research Trust | in-house | EW0000029 | sendai-virus |
| 110 | CL110.4 | Rett Syndrome Research Trust | in-house | EW0000029 | sendai-virus |

| | | | | | |
|---|---|---|---|---|---|
| 112 | CL112.10 | Rett Syndrome Research Trust | in-house | EW0000018 | sendai-virus |
| 112 | CL112.14 | Rett Syndrome Research Trust | in-house | EW0000018 | sendai-virus |
| 112 | CL112.15 | Rett Syndrome Research Trust | in-house | EW0000018 | sendai-virus |
| 113 | CL113.6 | Rett Syndrome Research Trust | in-house | EW0000026 | sendai-virus |
| 113 | CL113.8 | Rett Syndrome Research Trust | in-house | EW0000026 | sendai-virus |
| 114 | CL114.2 | Rett Syndrome Research Trust | in-house | EW0000036 | sendai-virus |
| 114 | CL114.5 | Rett Syndrome Research Trust | in-house | EW0000036 | sendai-virus |
| 115 | CL115.3 | Rett Syndrome Research Trust | in-house | EW0000042 | sendai-virus |
| 115 | CL115.8 | Rett Syndrome Research Trust | in-house | EW0000042 | sendai-virus |
| 116 | CL116.10 | Rett Syndrome Research Trust | in-house | EW0000051 | sendai-virus |
| 116 | CL116.13 | Rett Syndrome Research Trust | in-house | EW0000051 | sendai-virus |
| 116 | CL116.14 | Rett Syndrome Research Trust | in-house | EW0000051 | sendai-virus |
| 116 | CL116.6 | Rett Syndrome Research Trust | in-house | EW0000051 | sendai-virus |
| 116 | CL116.8 | Rett Syndrome Research Trust | in-house | EW0000051 | sendai-virus |
| 119 | CL119.2 | Rett Syndrome Research Trust | in-house | EW0000075 | sendai-virus |
| 119 | CL119.6 | Rett Syndrome Research Trust | in-house | EW0000075 | sendai-virus |
| 121 | CL121.1 | Rett Syndrome Research Trust | in-house | EW0000065 | sendai-virus |
| 121 | CL121.3 | Rett Syndrome Research Trust | in-house | EW0000065 | sendai-virus |
| 125 | CL125.20 | Rett Syndrome Research Trust | in-house | EW0000016 | sendai-virus |
| 125 | CL125.9 | Rett Syndrome Research Trust | in-house | EW0000016 | sendai-virus |
| 127 | CL127.1 | Rett Syndrome Research Trust | in-house | EW0000031 | sendai-virus |
| 127 | CL127.6 | Rett Syndrome Research Trust | in-house | EW0000031 | sendai-virus |
| 139 | CL139.5 | Rett Syndrome Research Trust | in-house | EW0000164 | sendai-virus |
| 139 | CL139.8 | Rett Syndrome Research Trust | in-house | EW0000164 | sendai-virus |
| 141 | CL141.3 | Rett Syndrome Research Trust | in-house | EW0000153 | sendai-virus |
| 141 | CL141.4 | Rett Syndrome Research Trust | in-house | EW0000153 | sendai-virus |
| 199 | CL199.5 | Coriell | Coriell | GM24666 | retrovirus |
| 200 | CL200.1 | Coriell | Coriell | AG25367 | sendai-virus |
| 201 | CL201.1 | Coriell | Coriell | GM24675 | retrovirus |

#### DETAILED MACHINE LEARNING METHODS

##### Feature extractor for brightfield morphology images

Brightfield embeddings were obtained using a ResNet18 convolutional neural network architecture to map images to 512-dimensional space. The model was trained in PyTorch using the Adam optimizer (default parameters) and arcloss with the following parameters: (s=32, m=0.5, easy_margin=True). At train-time the following data augmentations were applied: affine transforms, blur, brightness and contrast jittering, and deletion of image parts. Images from 22,000 organoids were used during training; ages of images from each organoid vary by less than a factor of 2.

To predict qPCR values from learned brightfield embeddings, RidgeRegression with default parameters provided by scikit-learn was used.

##### Feature extractor for DAPI-stained tissue (embedding of patch)

A ResNet-like fully convolutional model was used to embed patches into 128-dimensional space (PyTorch model definition below):

```
nn.Sequential(
        nn.Conv2d(  1,  64, stride=2, kernel_size=(5, 5), padding=2),
        nn.ELU(inplace=True),
        Residual(
                nn.Conv2d( 64,  64, stride=1, kernel_size=(3, 3), padding=1),
                nn.ELU(inplace=True),
                nn.Conv2d( 64,  64, stride=1, kernel_size=(3, 3), padding=1),
        ),
        Residual(
                nn.Conv2d( 64,  64, stride=1, kernel_size=(3, 3), padding=1),
                nn.ELU(inplace=True),
                nn.Conv2d( 64,  64, stride=1, kernel_size=(3, 3), padding=1),
        ),
        Residual(
                nn.Conv2d( 64,  64, stride=1, kernel_size=(3, 3), padding=1),
                nn.ELU(inplace=True),
                nn.Conv2d( 64,  64, stride=1, kernel_size=(3, 3), padding=1),
        ),
        nn.Conv2d( 64, 128, stride=2, kernel_size=(3, 3), padding=1),
        nn.ELU(inplace=True),
        LocalNormalization(),
        Residual(
                nn.Conv2d(128, 128, stride=1, kernel_size=(3, 3), padding=1),
                nn.ELU(inplace=True),
                nn.Conv2d(128, 128, stride=1, kernel_size=(1, 1), padding=0),
        ),
        Residual(
                nn.Conv2d(128, 128, stride=1, kernel_size=(3, 3), padding=1),
                nn.ELU(inplace=True),
                nn.Conv2d(128, 128, stride=1, kernel_size=(1, 1), padding=0),
        ),
        nn.Conv2d(128, 128, stride=2, kernel_size=(3, 3), padding=1),
```

```
        LocalNormalization(),
        Residual(
                nn.Conv2d(128, 128, stride=1, kernel_size=(3, 3), padding=1),
                nn.ELU(inplace=True),
                nn.Conv2d(128, 128, stride=1, kernel_size=(1, 1), padding=0),
        ),
        LocalNormalization(),
)
```

Max pooling is not used, which allows the model to track sizes and distances precisely. LocalNormalization corresponds to position-wise L2 normalization of embedding. The model was trained in PyTorch using the Adam optimizer (default parameters, except learning rate 1e-4) and a new custom loss function that builds upon a contrastive loss approach. The key modification is allowing a limited number of distinct groups (dubbed “clusters”) within a single region. Loss is only computed for patches with significant amounts of DAPI fluorescence. Training is done in mixed precision (fp16+fp32) provided by Apex (opt_level="O1"). At train-time, the following data augmentations were used: affine transformations (rotations and shifts, minimal scaling), brightness and contrast jittering. Images from 90 organoids were used during training. Images had either multiple z planes or images taken at different timepoints.

**Classification model to predict disease state from patch**

RandomForest provided by scikit-learn with following parameters

```
RandomForestClassifier(min_samples_leaf=30, n_estimators=30, class_weight='balanced',
random_state=12)
```
